# A novel chitinase-like family of candidate effectors unique to aphids

**DOI:** 10.1101/2025.08.07.665532

**Authors:** Rosa Lopez-Cobollo, Simone Altmann, Peter Thorpe, Nadine Douglas, Po-Yuan Shih, Laura Eccleston, Mark Lord, Eunji Hong, Emese Klug, Javaid Iqbal, Jorunn Bos, James Carolan, Colin Turnbull

**Affiliations:** Department of Life Sciences, Imperial College London, London SW7 2AZ, UK; Division of Plant Sciences, School of Life Sciences, University of Dundee, Dundee, DD5 4EH, UK; Division of Computational Biology, School of Life Sciences, University of Dundee, Dundee, DD5 4EH; Department of Biology, Maynooth University, Republic of Ireland; School of Biology and Environmental Science, University College Dublin, Dublin 2, Republic of Ireland; The James Hutton Institute, Invergowrie, Dundee DD2 5DA, UK

## Abstract

Molecular interactions between aphids and plants include delivery of salivary effector proteins into host cells, acting as virulence factors to suppress host immunity, or as avirulence functions triggering immune activation. However, understanding of virulence and avirulence mechanisms in aphid-plant systems is currently limited. Here, we report discovery of an effector candidate family that is unique to aphids. Using functional genomics data on divergent pea aphid (*Acyrthosiphon pisum*) genotypes and their F1 progeny, we filtered for differentially expressed saliva proteins that co-segregated with virulence or avirulence phenotypes. LOC100575698 (ACPISUM_029930), annotated as an uncharacterized protein, was the sole candidate effector for which RNA-Seq and saliva proteomics data showed significantly different expression both between avirulent and virulent parents and between their segregating F1 progeny, with this gene upregulated in avirulent genotypes. BLASTP searches revealed multiple divergent homologs only in genomes of the Aphidomorpha infra-order, suggesting a hitherto undefined ancient aphid-specific gene family. AlphaFold models indicate strong structural similarities but weak sequence homology to chitinases. Because the aphid-specific clade all lack canonical DxxDxDxE motifs for catalytic activity, we designate the proteins as a novel CHitinase-Like (CHL) family. Association of ACPISUM_029930 (ApCHL1) with avirulence was further supported by co-segregating SNPs and a genotype-specific alternatively spliced isoform. We hypothesise that CHL proteins may function similarly to phylogenetically unrelated chitin-binding fungal effectors that sequester chitin, also present in aphid stylets, potentially preventing defence activation through plant chitin receptors and/or blocking chitin degradation by host-secreted chitinases.

## INTRODUCTION

Aphids are global plant parasites that cause widespread damage to crops through their sap sucking and through acting as vectors for many plant viruses. In agriculture, aphids are managed through insecticides, host resistance genes or biocontrol agents. None of these strategies is fully effective, with aphids often evolving to overcome agrochemicals or resistance genes. In addition to interest in fundamental mechanisms of aphid-plant interactions, there is therefore a strong motivation to develop novel robust and sustainable routes to aphid control.

A few plant resistance genes, mostly encoded by coiled-coil, nucleotide binding, leucine-rich repeat (abbreviated to CC-NLR or CNL) immune receptors, that are highly effective against aphids have been discovered and deployed in crops such as tomato (*Mi-1*) and melon (*Vat*), along with further examples in rice that act against related phloem-feeding brown planthopper (BPH) pest species (Dogimont et al., 2014; Du et al., 2009; Rossi et al., 1998).

Aphids deliver a range of effectors via saliva into host plant tissues, generally acting as virulence factors for example by suppressing plant defences. Effector annotations in many cases point to putative, but as yet mostly unproven, functions such as cell-wall degrading cellulases and pectinase (van Bel and Will, 2016). In addition, there are now several studies directly studying aphid effector functions. Some of these have provided strong evidence for interactions with putative host targets such as Me10 withTFT7 (Chaudhary et al., 2019), and Mp1 with VPS52 (Rodriguez et al., 2017). Others have expressed effectors *in planta* to test for subsequent effects on aphids, with quite diverse results from increased to decreased virulence performance (Bos et al., 2010; Elzinga et al., 2014; Pitino and Hogenhout, 2013). Knockdown of effectors upon aphid feeding on plants expressing RNAi sequences in several studies has indicated impacts on aphid performance (Yu et al., 2016). There are also insights on how aphid-detecting NLRs, such as the sensor NLR Mi-1.2, act with cell surface proteins and the helper NLR NRC4 (Peng et al., 2016; Wu et al., 2017). Notably though, none of these interactions have yet tracked all the way to how specific aphid effectors may enable effector-triggered immunity (t) via canonical activation of R genes. Recently, the aphid Mp10 effector, a member of the chemosensory protein (CSP) family was shown to interact directly with an AMSH deubiquitinase host target protein (Gravino et al., 2024). This interaction has negative consequences for PAMP-triggered immunity (PTI) acting via cell surface receptors such as FLS2 and EFR, and is dependent on the BAK1 co-receptor. Interestingly, AMSH proteins may also be involved in ETI (Schultz-Larsen et al., 2018) and Mp10 may also interact with ETI receptors such as the TIR domain NLR (TNL) protein RCSP (Rao et al., 2024). In BPH, the BISP effector acts in dual modes in host cells, both suppressing PTI via interaction with a receptor-like cytoplasmic kinase and triggering ETI via the CNL BISP14 (Guo et al., 2023). Despite these several recent breakthroughs, the majority of aphid effectors remains uncharacterised and their modes of action in host plant tissues are largely unknown.

To address the knowledge gaps on aphid effectors, an initial research goal has been to generate comprehensive catalogs of candidate effectors present in salivary proteomes, with data now being available across multiple aphid species (Carolan et al., 2011, 2009; Chaudhary et al., 2015, 2014; Thorpe et al., 2024, 2016; Waksman et al., 2024; Zhang et al., 2021). Such datasets can be described as effectoromes as they represent the complement of proteins potentially delivered into host plants. In almost all cases, these studies have employed artificial diet approaches to collect salivary samples, mainly because of the difficulty of detecting the minute quantities of saliva proteins diluted into the entire plant proteome of aphid infested tissues. It remains possible that the saliva proteome composition changes qualitatively and/or quantitatively following aphid interactions with their hosts. One recent study detected aphid proteins from bamboo tissue on which aphids had previously fed (Zhang et al., 2023). In other studies, salivary gland proteomes and transcriptomes have enabled prediction of likely salivary effectors, with data sets typically filtered for proteins carrying secretion signals (Carolan et al., 2011), and in some cases also for proteins lacking trans-membrane domains (Pavithran et al., 2024; Zhang et al., 2023).

We recently extended the depth of data on pea aphid candidate effectors through transcriptomics in parallel with proteomics of saliva and salivary glands (Thorpe et al., 2024). In this study, we compared transcripts and proteins of aphid genotypes with highly contrasted virulence (VIR) and avirulence (AVR) phenotypes on *Medicago truncatula* host plants carrying the *RAP1* R gene QTL (Kanvil et al., 2015; Stewart et al., 2009). Through examination of F1 hybrid populations, we uncovered subsets of transcripts and proteins whose upregulation co-segregated with VIR or AVR phenotypes (Thorpe et al., 2024). There were 47 saliva proteins with differential abundance between the VIR and AVR parents, many of which were also differentiated in transcriptomes and salivary gland proteomes. This protein set was dominated by exopeptidases (40%) and uncharacterised proteins (23%). Here we focus on the single gene (ACPISUM_029930; uncharacterized protein LOC100575698) that was differential in all comparisons: parental transcriptomes, F1 transcriptomes and parental saliva proteomes. We extend the data to reveal that this protein co-inherits with avirulence in saliva of F1 aphids. Moreover, the protein is a member of a novel chitinase-like (CHL) family exclusive to aphids, for which multiple members are present at high abundance in saliva.

## RESULTS

### A single salivary effector protein co-segregates with pea aphid avirulence phenotypes

We previously found several salivary proteins that were differentially abundant between parental aphid genotypes N116 and PS01 that display virulent (VIR) and avirulent (AVR) phenotypes, respectively, on *M. truncatula* host plants carrying the *RAP1* resistance QTL (Thorpe et al., 2024). Some of the underlying genes were similarly differentially expressed in RNA-Seq experiments both on parental and on F1 segregating populations. Here we tested whether the differential patterns are also expressed in salivary proteomes of representative F1 virulent and avirulent individuals from the same populations. Across all the saliva samples analysed, a total of 123 proteins were confidently identified, with 36 proteins common to all genotypes (Supplementary Fig. S1; Supplementary Table S1). We were particularly interested in proteins that were more abundant in, or unique to, both parental VIR and F1 VIR aphids compared with the corresponding AVR samples, and vice-versa. No such proteins were found only in VIR genotypes. However, there was a single protein unique only to both the parental AVR and the F1 AVR clones AVR data: ACPISUM_029930 (uncharacterized protein LOC100575698).

To analyse the differential expression of ACPISUM_029930 in more detail, we first extracted the published RNA-Seq data (Thorpe et al., 2024) for both the parental AVR and VIR aphids, and for the AVR and VIR F1 pools. These data indicate ∼nine-fold higher expression in avirulent clones (*P* = 3.5x10^-6^, 1.5x10^-6^ by One-Way ANOVA for parents and F1, respectively) (Fig. 1A). Because the F1 aphids were analysed originally as bulk segregant pools, we next measured ACPISUM_029930 expression by-qRT-PCR for both parental lines, and for selected individual progeny, three each from virulent and avirulent members of the F1 populations. These analyses confirmed that avirulent clones all had significantly higher expression than virulent clones on A17 host plants that carry the *RAP1* aphid resistance QTL (eight-fold; overall *P* = 5.3x10^-7^, Fig. 1B). ACPISUM_029930 was also upregulated in the same clones on the susceptible DZA315.16 host genotype (four- to seven-fold; Supplementary Fig. S2) and on the universally compatible legume *V. faba* (three-fold), indicating that increased expression was not solely due to interaction with an incompatible host. Although there was no significant difference in expression on *M. truncatula* genotypes carrying or lacking *RAP1*, ACPISUM_029930 was up to two-fold more highly expressed in some AVR clones on *M. truncatula* compared with those on *Vicia faba* (P<0.05).

**Fig. 1.**
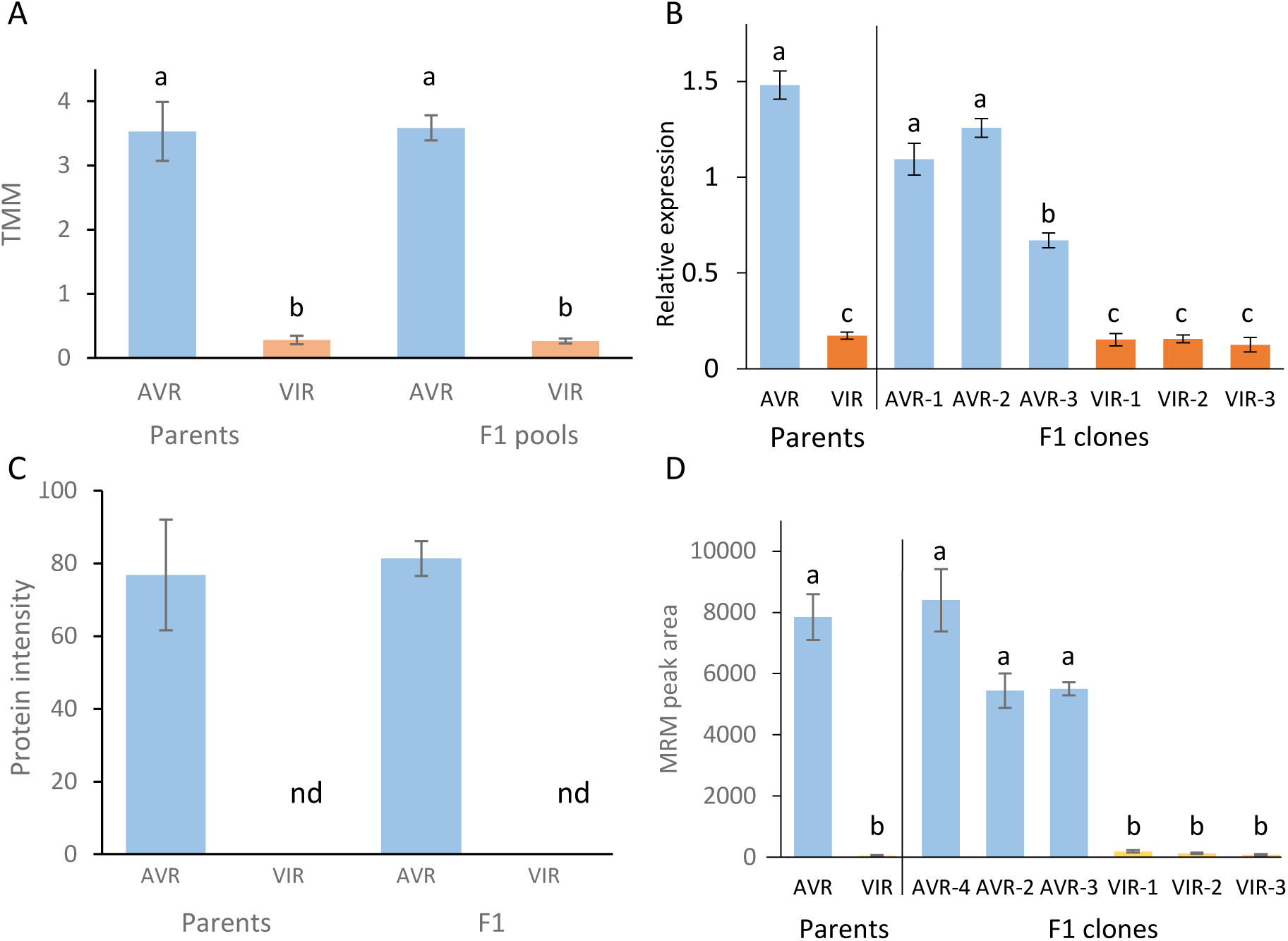
Co-segregation of ACPISUM_029930 differential expression with avirulence phenotypes at both RNA and protein levels. A. Expression of ACPISUM_029930, measured by RNA-Seq. Aphids collected 24 h after infestation on resistant near-isogenic line (RNIL) plants carrying *RAP1.* F1 samples were from 20 pooled clones each of avirulent and virulent phenotypes. Data are Transcripts per Million Mapped reads. N = 5 independent libraries for each sample type. B. Expression of ACPISUM_029930, analysed by qRT-PCR. Aphids collected 24 h after infestation on A17 plants. Genotypes were AVR and VIR parents, and three representatives each of AVR and VIR F1 progeny. Values are expression relative to mean of reference genes GAPDH and SDHB. N = 3 independent biological replicates. C. ACPISUM_029930 protein level in saliva, analysed by accurate mass LC-MS. Genotypes were AVR and VIR parents, and one representative each of AVR and VIR F1 progeny. Data are relative intensities measured by label-free quantitation. N = 3 independent biological replicates. nd, not detected. D. ACPISUM_029930 protein level in saliva, measured by Multiple Reaction Monitoring LC-MS. Genotypes were AVR and VIR parents, and three representatives each of AVR and VIR F1 progeny. MRM peptide was YIDTIDPTAK.+2y8. N = 3 independent biological replicates. Plots are means ± SE. For A, B and D, letters in common above bars indicate values that are not significantly different by one-way ANOVA and Tukey’s HSD test.

We then focussed on quantitative analysis at the protein level, using saliva samples collected via artificial diets, as that would give indications of relative amounts of effectors delivered into host tissues. Two sets of analyses again pointed strongly to co-segregation of differential expression at the protein level, with patterns that closely match the transcriptional data. Although several salivary proteins were differentially abundant or unique between parental aphid clones (Thorpe et al., 2024),

ACPISUM_029930 was again unique in also being differentially abundant in saliva between the F1 virulent and avirulent progeny. First, we used untargeted LCMS analysis that indicated presence of ACPISUM_029930 both in the AVR parent and a representative avirulent F1 clone (AVR-1), but not in the VIR parent nor in a virulent F1 clone (VIR-1) (Fig. 1C). To expand on this finding, we undertook targeted analysis of ACPISUM_029930 via LCMS-with Multiple Reaction Monitoring (MRM) for both parents and three F1 clones of each phenotype. These accurate quantifications confirmed the presence of ACPISUM_029930 in all AVR saliva samples, whereas the protein was near or below the detection limit for all VIR samples (Fig. 1D). Quantifying the peak abundance for the strongest MRM peptide signal (YIDTIDPTAK.+2y8) gave an estimated 60-fold higher level of ACPISUM_029930 protein across all avirulent clones, and protein identity was confirmed from a second MRM peptide (EIQAQGIVTK.+2y7; Supplementary Table S2).

### The candidate effector ACPISUM_029930 is alternatively spliced, with differential isoform abundance between avirulent and virulent genotypes

The gene model corresponding to ACPISUM_029930 at LOC100575698 on published genome assemblies at Aphidbase and NCBI predicts a gene with 7 exons, encoding a protein of 258 amino acids. However, mapping of our RNA-Seq data to these genomes indicates two alternatively spliced variants (Fig. 2A; Supplementary Fig. S3), with the short form corresponding to the published gene models. The longer form (ACPISUM_029930-T1) has 8 exons and encodes a protein of 371 amino acids. Predicted protein sequences are given at Supplementary Table S3. Further examination of the mapped RNA-Seq reads indicates that there is also differential abundance of the two isoforms. The long isoform was undetectable in both the VIR parent and the pooled VIR subset of the F1 population, with all reads mapping to the short isoform. In contrast, both isoforms were expressed in the avirulent parent PS01 and only the long isoform in the pooled AVR F1 samples (Supplementary Fig. S3). Sufficient reads were detected from PS01 samples to calculate that the expression of the long isoform was about three-fold greater than the short isoform (Fig. 2B). Based on SNP analysis, PS01 is heterozygous at this locus, with only one allele being expressed as the long form. However, the short form was able to be expressed off both PS01 alleles (Table 1; Supplementary Table S4). Analysis of the LCMS tryptic peptide sequences from ACPISUM_029930 revealed one peptide mapping exclusively to the long isoform, spanning exons 2 and 4, and multiple peptides for the shared protein region (Supplementary Table S5). Taken together, the evidence strongly supports a model where ACPISUM_029930 is predominantly expressed as a long isoform encoded by one allele present in the AVR parent. Moreover, this allele is inherited exclusively by individuals with an AVR phenotype in F1 populations.

**Fig. 2.**
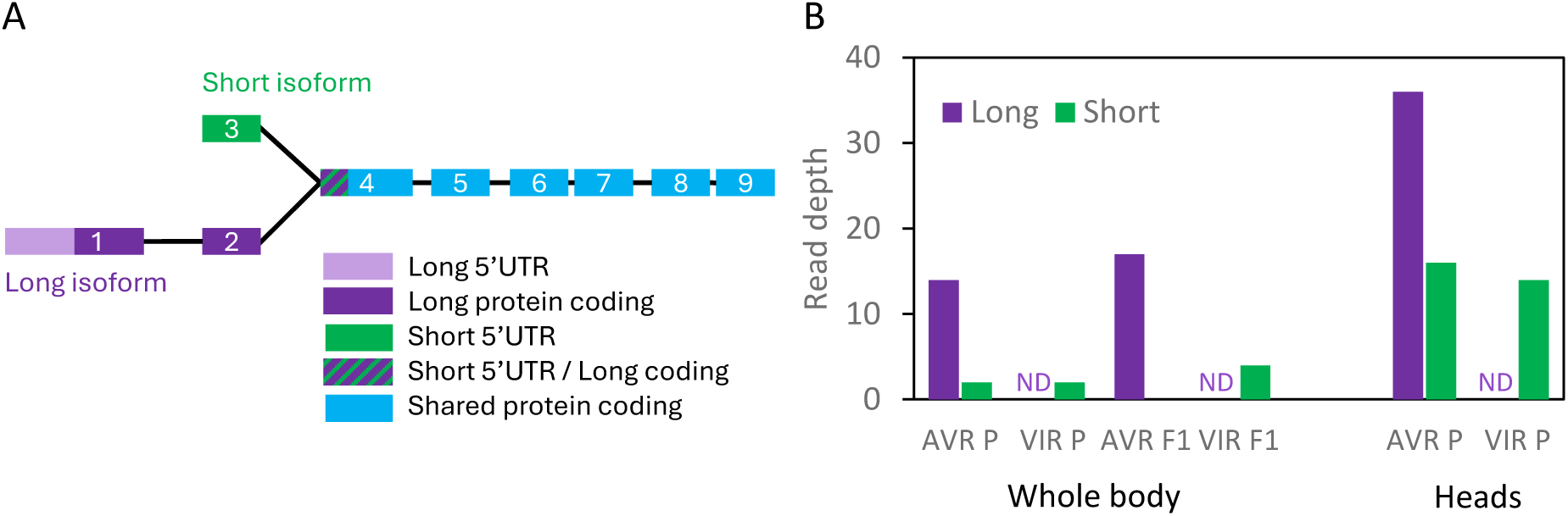
Genotype-specific expression two splice isoforms of ACPISUM_029930. The long and short isoforms encode proteins of 371 and 258 amino acids, respectively. A, schematic of exon - intron structure. Diagram is not to scale. B, the long isoform is expressed only in AVR genotypes. Data are summed read depths, extracted from RNA-Seq tracks. Samples were from AVR and VIR parents (P) and from pooled F1 AVR or VIR progeny, as reported in (Thorpe et al., 2024).ND, not detected.

**Table 1.** Differential expression of ACPISUM_029330 isoforms. SNPs in exon 1 and exon 4 enabled assignment of RNA-Seq reads to long and short isoforms. Data are read counts. Nil = no reads detected in this location.

| Genotype | Exon 1 |  | Intron spanning | Exon 4 |  | Intron spanning | Intron spanning |
| --- | --- | --- | --- | --- | --- | --- | --- |
|  | C (Ref) | A (Alt) | Exon 1 to 2 (Long) | A (Ref) | G (Alt) | Exon 2 to 4 (Long) | Exon 3 to 4 (Short) |
| AVR parent (PS01) | 0 | 23 | 17 (A) | 5 | 32 | 22 (G) | 2 (A), 2 (G) |
| VIR parent (N116) | Nil | Nil | Nil | 10 | 0 | Nil | 8 (A) |
| AVR F1 pool | 0 | 11 | 9 (A) | 0 | 14 | 12 (G) | nil |
| VIR F1 pool | Nil | Nil | Nil | 4 | 0 | Nil | 4 (A) |

### ACPISUM_029930 is a member of an aphid-specific chitinase-like (CHL) family with distant homology to true chitinases

Because ACPISUM_029930 (LOC100575698) is annotated as an uncharacterised protein in public databases, we applied a range of sequence-based and structure-based bioinformatic approaches to attempt to reveal its putative function. BLASTP searches of the pea aphid genome at NCBI revealed seven genes, using a cutoff of E<10^-5^ (Table 2). The same BLASTP query without taxonomic restriction returned approximately 200 BLASTP hits, all to aphid species, with only two, both from *Cinara cedri*, having functional annotations as Glycoside hydrolase superfamily (Supplementary Table S6; Supplementary Fig. S4). A phylogenetic tree using sequences from a representative subset of aphid species is shown in Figure 3. The significant homologies strongly support existence of a hitherto unrecognised gene family. Despite the low overall sequence identities across these proteins, multiple sequence alignment revealed several highly conserved amino acids (Fig. 5G).

**Fig. 3.**
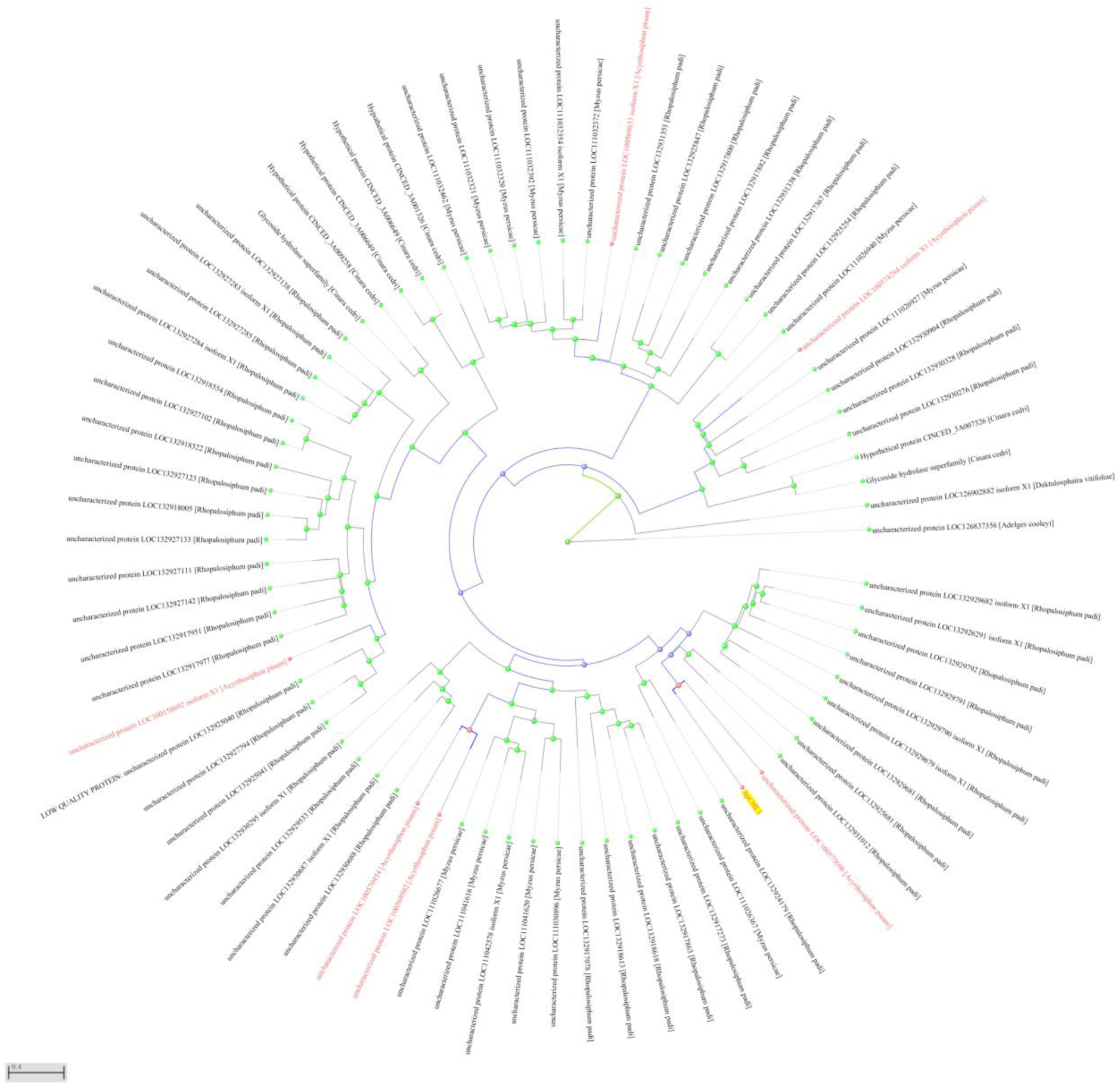
ACPISUM_029930 is a member of a previously unknown gene gamily, found only in aphids. Unrooted Neighbour Joining tree based on BLASTP hits to ACPISUM_029930 protein sequence, with E < 10^-5^ cutoff. Search was restricted to representative species within the infra-order Aphidomorpha: *A. pisum*, *Myzus persicae*, *Rhopalosiphum padi*, *Cinara cedri*, *Daktulosphaira vitifoliae* and *Adelges cooleyi*. Isoforms were removed, as were short predicted proteins <250 amino acids. An equivalent tree for BLASTP to all species is shown at Supplementary Fig. S4.

**Table 2.** ACPISUM_029930 has multiple paralogs in the pea aphid genome.

| CHL gene | LOC | NCBI protein | ACPISUM gene | ACYPI gene | Predicted protein length (aa) |
| --- | --- | --- | --- | --- | --- |
| CHL1 | LOC100575698 | XP_016660502.1* | ACPISUM_022930 | ACYPI53036 | 371 (258*) |
| CHL2 | LOC100570454 | XP_003242981.1 | ACPISUM_008664 | ACYPI28420 | 342 |
| CHL3 | LOC100569502 | XP_016658069.2 | ACPISUM_029311 | ACYPI28420 | 306 |
| CHL4 | LOC100574284 | XP_003246661.1 | ACPISUM_019736 | ACYPI49582 | 358 |
| CHL5 | LOC100158692 | XP_016659499.1 | ACPISUM_012348 | ACYPI000132 | 368 |
| CHL6 | LOC100569633 | XP_003246843.3 | ACPISUM_020227 | ACYPI54505 | 354 |
| CHL7 | LOC100167587 | XP_029347624.1 | ACPISUM_020227 | ACYPI54505 | 276 (686**) |

\*short isoform; \*\*expression data do not support this gene model

We next used PSI-BLAST (NCBI) across all taxa to seek additional, weaker sequence-based homologies that might uncover more distantly related proteins. Searching against genomes for Sternorrhyncha, the sub-order within Hemiptera that includes aphids and several other plant feeding groups, returned >400 hits (Supplementary Table S7). Phylogenetic trees constructed from the protein sequences indicated sub-clades that included many proteins annotated as chitinases, Imaginal disc growth factors (IDGFs), mucins or flocculins (Supplementary Fig. S5). Species represented in these clusters included aphids and also non-aphids such as whitefly (*Bemisia tabaci*) psyllid (*Diaphorina citri*) and mealybug (*Phenacoccus solenopsis*). Importantly, however, these groupings are all distinct from the aphid-specific clade that contains ACPISUM_029930. Notably, none of the proteins in the aphid-specific sub-clade contain the canonical DxxDxDxE motif necessary for chitinase catalytic activity (Supplementary Fig. S6; Supplementary Table S8), annotated as the active site of glycosyl hydrolase family 18 proteins at InterPro (IPR001579). We conclude that this group of proteins is distantly but significantly related to chitinases and related proteins that are present across all insect groups. We therefore name the aphid-specific proteins as a novel CHitinase-Like (CHL) family, with ACPISUM_029930 being the founding member, designated as ApCHL1. Because the large number of hits hindered visualisation, we re-submitted the ApCHL1 (ACPISUM_029930) PSI-BLAST query instead with taxonomic restriction to *A. pisum* and three additional members of Aphidomorpha: *Myzus persicae*, *D. vitifoliae* and *A. cooleyi*, along with other related Hemiptera: the psyllid *D. citri* and the whitefly *B. tabaci*, and a distant insect relative, *Drosophila melanogaster*, aiming to reveal the distribution of genes across insect groups (Fig. 4; Supplementary Table S9). This analysis confirmed the aphid-specific clade, whereas chitinases, IDGF and mucins appear to be widely represented across all insects.

**Fig. 4.**
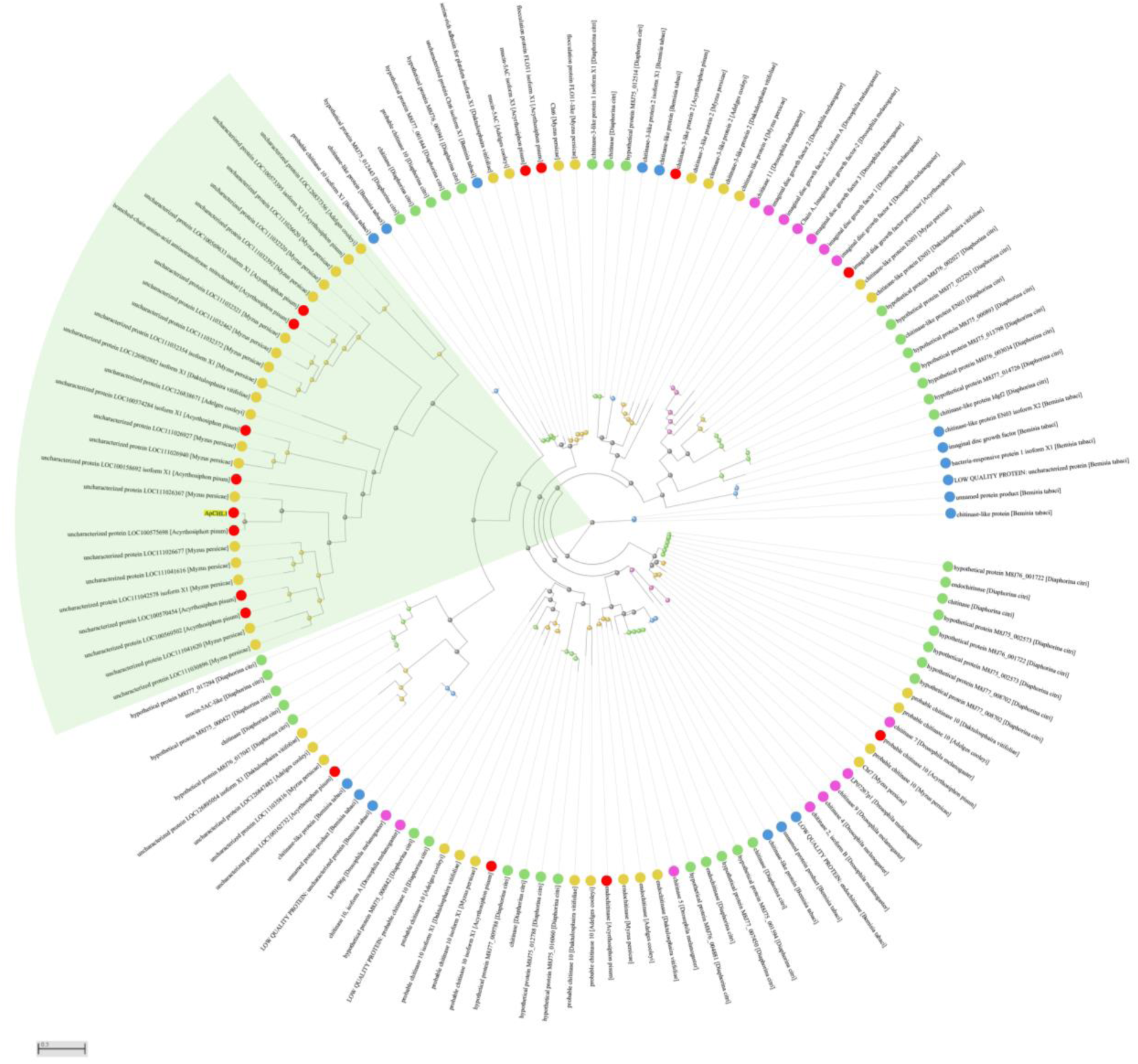
The aphid-specific group of proteins is distantly related to chitinases and other proteins present in aphids and all other insect groups. Protein set was returned after four rounds of PSI-Blast to ApCHL1, with taxonomic restriction to *A. pisum* (red circles), three additional species within Aphidomorpha (gold): *Myzus persicae*, *Daktulosphaira vitifoliae* and *Adelges cooleyi*; other closely related Hemiptera: *Diaphorina citri* (psyllid, green) and *Bemisia tabaci* (whitefly, blue); and a distant insect relative, *D. melanogaster* (magenta). Isoforms and short predicted proteins (<250 amino acids) were removed before creating an unrooted Neighbour Joining tree. The aphid-specific CHL clade is marked by a green segment. The input proteins are listed at Supplementary Table S9

Numbers of *CHL* genes varied greatly across species within the true aphids (Aphididae), from two in the Russian wheat aphid (*Diuraphis noxia*) to 44 in the Bird cherry-oat aphid (*Rhopalosiphum padi*) (Table 3). In sister species with available genome data within the wider infraorder Aphidomorpha, *Adelges cooleyi* has two *CHLs*, whereas only a single *CHL* appears to be present in the phylloxeran *Daktulosphaira vitifolia.* Scrutiny of genomes for other sap-sucking species within the order Hemiptera confirmed that no *CHLs* are present outside the Aphidomorpha.

**Table 3.** Number of chitinase-like (CHL) genes in genomes of aphids and other selected Hemiptera relatives. Genes were retrieved from NCBI by BLASTP against ApCHL1, Cutoff E = 10^-5^. Where isoforms were annotated, only the longest protein sequence was retained. Predicted proteins <250 or >650 amino acids were excluded. A full phylogenetic tree for CHL proteins in aphids and other Aphidomorpha species is shown at Supplementary Fig. S4.

| Species | Number of CHL genes |
| --- | --- |
| **Aphids** |  |
| *A. pisum* | 6 (7) |
| *Aphis craccivora* | 6 |
| *Aphis glycines* | 11 |
| *Aphis gossypii* | 16 |
| *Cinara cedri* | 7 |
| *Diuraphis noxia* | 2 |
| *Macrosiphum euphorbiae* | 11 |
| *Melanaphis sacchari* | 15 |
| *Metopolophium dirhodum* | 13 |
| *Myzus persicae* | 14 |
| *Rhopalosiphum maidis* | 21 |
| *Rhopalosiphum padi* | 44 |
| *Semiaphis heraclei* | 5 |
| *Sipha flava* | 4 |
| *Uroleucon formosanum* | 23 |
| **Sister species within Aphidomorpha** |  |
| *Adelges cooleyi* | 2 |
| *Daktulosphaira vitifolia* | 1 |
| **Sister species in wider Sternorrhyncha** |  |
| *Bemisia tabaci* | 0 |
| *Diaphorina citri* | 0 |
| *Phenacoccus solenopsis* | 0 |
| **Sister species in wider Hemiptera** |  |
| *Laodelphax striatellus* | 0 |
| *Nilaparvata lugens* | 0 |

### CHL proteins have highly conserved structures, with similarities to chitinases

We next used AlphaFold to predict structures of CHL proteins. High confidence models were generated (Fig. 5A; ApCHL1 pTM = 0.93), indicating that CHLs adopt a TIM-barrel configuration. Submitting the predicted structure of ApCHL1 to Foldseek returned multiple significant hits. The closest structures were all from aphids, representing other CHLs (Supplementary Fig. S7; Supplementary Table S10). Notably, there was also strong similarity of ApCHL1 structure to chitinases such as the insect protein from *Ostrinia furnicalis* (Fig. 5C; PDB: 6JAX). SwissDock modelling of ApCHL1 with a chitin tetramer (CO4) ligand indicated that CHLs may have the potential to bind chitin oligomers (Fig. 5B). The modelled binding affinity was -11.4 kcal/mol. The predicted position of the CO4 ligand on ApCHL1 was similar but not identical to that of the *O. furnicalis* PDB structure that was solved in the presence of the octamer CO8 (Fig. 5D). Creating an overlay of six AlphaFold-predicted CHLs from *A. pisum* using PyMol indicated highly similar structures despite the low sequence homology (Fig. 5E). Nonetheless, a WebLogo representation of predicted CHL sequences indicated several highly conserved residues (Fig. 5G; Supplementary Fig. S8), of which cysteines were by far the most common. The conserved cysteines are predicted to form disulphide bridges (Fig. 5F) and likely contribute to CHL protein folding

**Fig. 5.**
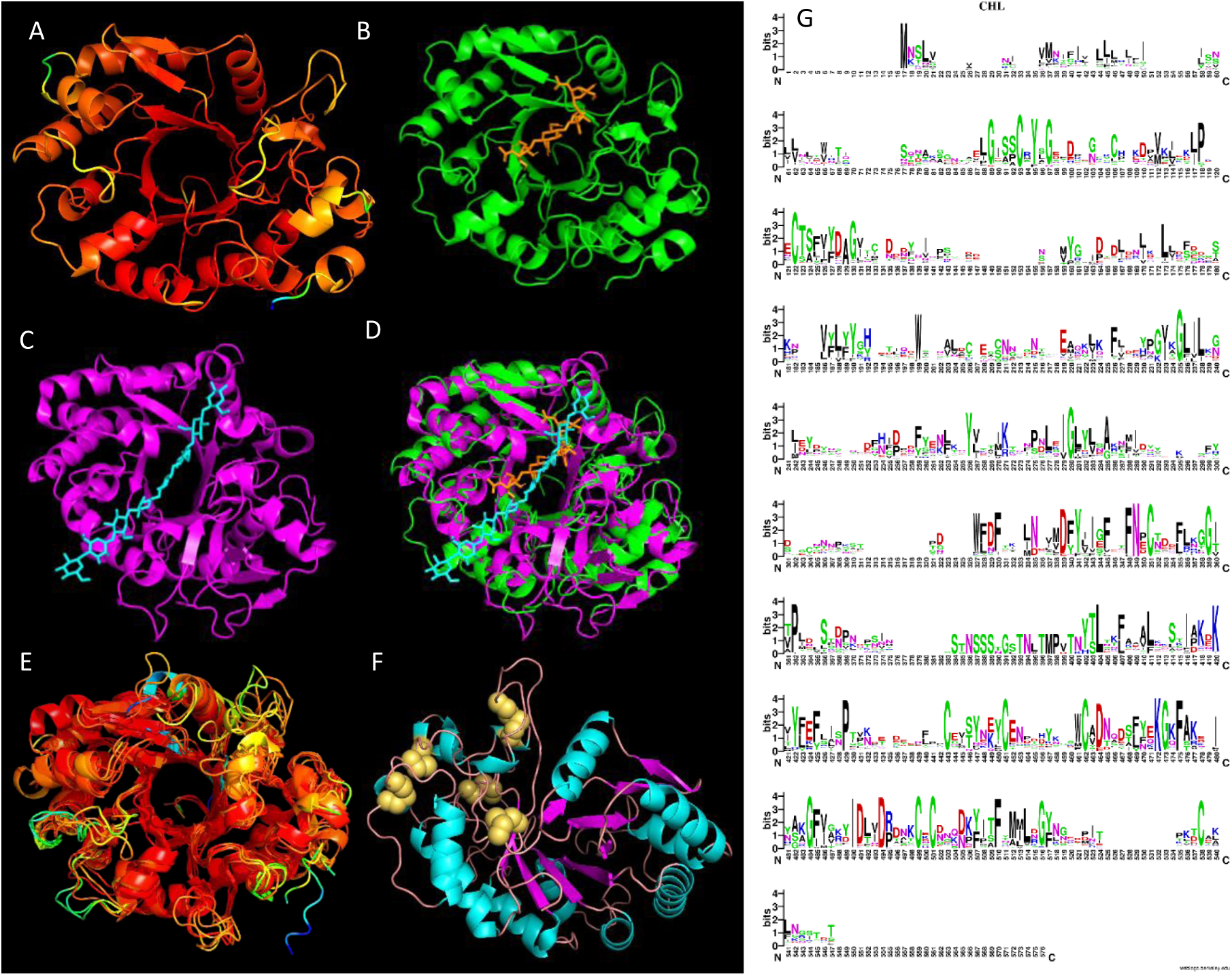
Structural models, ligand binding predictions and conserved residues of CHL proteins. A, AlphaFold model of ApCHL1 (pTM = 0.93). Rainbow coded for model confidence from red (high) to blue (low). B, SwissDock modelling of minimum energy configuration of chitin octamer (orange) binding to ApCHL1 (green). C, PDB model (6JAX; purple) of insect chitinase chain from *Ostrinia furnicalis*, in presence of chitin octamer (blue). D, Overlay of B and C. E, Overlay of Alphafold models for six *A. pisum* CHL proteins found in saliva. Average TM score 0.861; RMSD 1.76 Å; mean sequence identity 22%. F, ApCHL1 model rotated to show disulphide pairs (gold). G, WebLogo representation of aligned CHL proteins. Dataset is 60 proteins retrieved from BlastP of ApCHL1 at NCBI NR. All sequences are from aphids (family Aphididae). Isoforms and sequences <350 or >400 amino acids were removed prior to COBALT alignment. E-value cutoff 1e^-18^.

### Multiple CHL proteins are present in saliva of all studied aphid species

Following the discovery of multiple CHL genes in all available aphid genomes, we re-interrogated our previous saliva proteome data (Thorpe et al. 2024), which revealed that all predicted pea aphid CHL proteins were present in saliva. Moreover, label-free quantitative proteomics indicated that CHL proteins in pea aphid saliva represented four of the top six most abundant unannotated salivary proteins (Supplementary Table S1), pointing to likely functional importance and redundancy within the family.

Based on NCBI genome inspection and mapping of peptides detected by salivary proteomics, all seven CHLs predicted from BLASTP searches (Table 2) were found in *A. pisum* saliva (Table 3). Extraction of previously published RNA-Seq and saliva proteomics data for all ApCHLs indicated that ApCHL1 was the only family member that was significantly upregulated in both AVR parent and AVR F1 clones (Supplementary Table S11). Most of the CHL genes also have alternatively spliced isoforms, based on RNASeq exon/intron mapping. In all cases RNA-Seq read counts indicate a single dominant isoform. There are two pairs of tandem duplications, based on CHL2/CHL3 and CHL6/CHL7, with 89% and 71% amino acid sequence identity, respectively between each pair in their aligned regions. CHL7 on the AL4f genome at NCBI (LOC100167587) is called as a 686aa protein, and annotated as a branched chain amino acid aminotransferase (BCAT), but this may be an incorrect gene model. The N terminal ∼275 aa has CHL homology, whereas the C terminal portion has BCAT homology, and is much more strongly expressed (20x higher) than the CHL region. All the peptides detected by saliva proteomics map to the CHL sequence and none to BCAT, leading to the conclusion that the BCAT sequence is not expressed as a protein in saliva.

Available full length protein sequences were submitted to online tools (SignalP 6.0, OutCyte 2.0) for prediction of secretion. All detected ApCHL salivary proteins contain either an SP or a UPS predicted secretion motif (Table 3).

We then searched published saliva proteomes of other aphid species (*M. persicae*, *M. euphorbiae*, *R. padi*, *Sitobion avenae*). Multiple CHLs are present in saliva proteomes of all these aphid species (Table 4). Putative orthologs of ACPISUM_012348 and ACPISUM_008664 were present in saliva of all six species with detailed salivary proteomes available, and all species had at least four CHL proteins in saliva. The *M. persicae* ortholog of ACPISUM_008664 has previously been reported as Mp57, and had a significant negative effect on aphid performance when expressed in host plants (Elzinga et al., 2014). A recent computational study on *M. persicae* effectors (Waksman et al., 2024) included structural predictions of several proteins that we can now assign as CHLs, with their “Mp” gene numbers listed in Table 3. Two of the Mp CHLs (orthologs of ApCHL2 and ApCHL4) were reported to be under positive selection based on high dN/dS ratios (Thorpe et al., 2016).

**Table 4.**
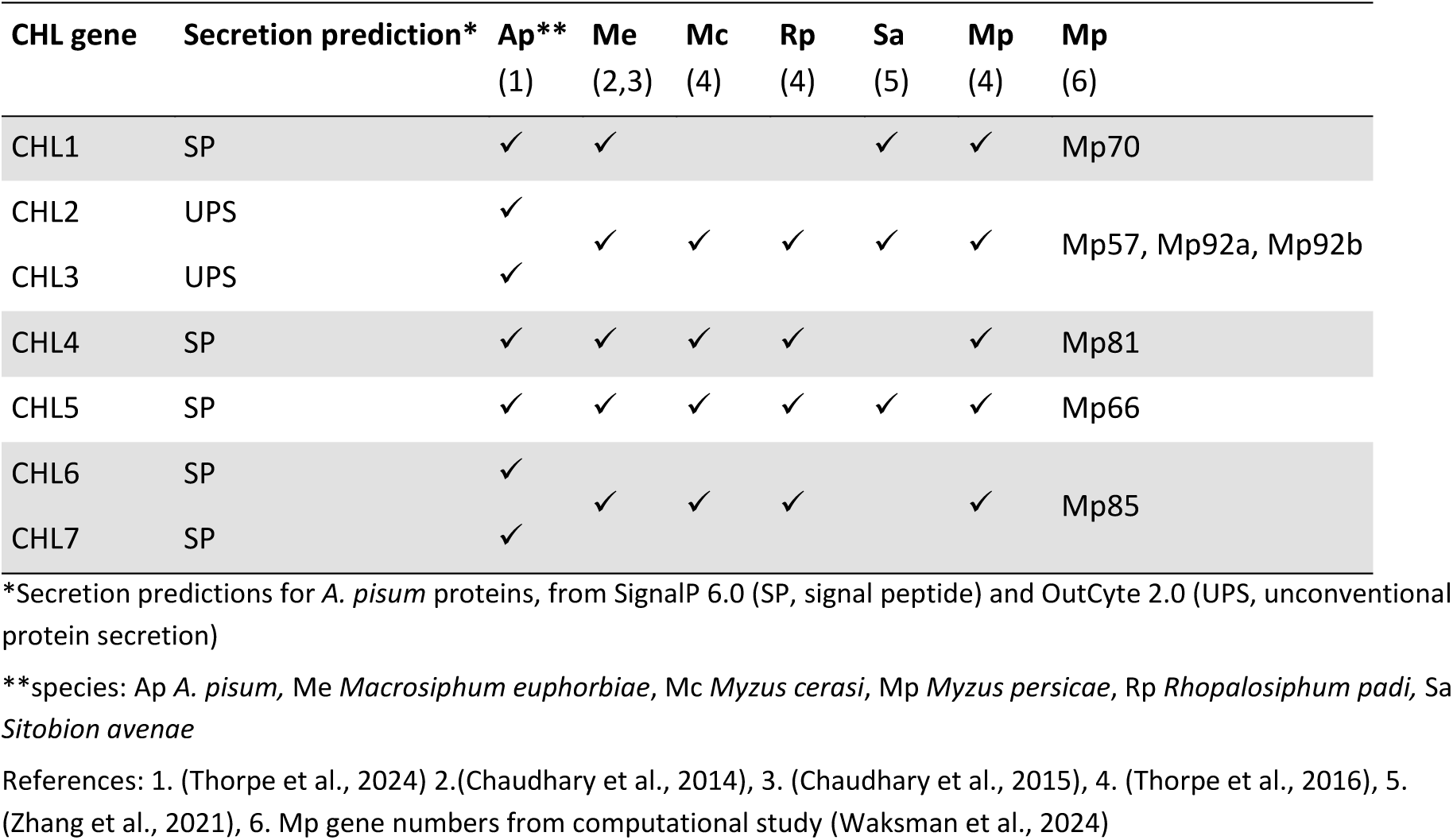
CHL homologs are common salivary proteins across aphid species. In pea aphid, there are two tandem duplicated gene pairs: CHL2/CHL3 and CHL6/CHL7. Therefore, where precise orthology relationships are unclear, CHL proteins from other aphids have been assigned to more than one pea aphid protein. ✓, reported in saliva.

## Discussion

### Co-segregation of ApCHL1 expression and avirulence phenotype

A combination of genetic and genomic approaches led to the discovery of the candidate effector gene ACPISUM_029930/LOC100575698, an uncharacterised gene that we have designated as ApCHL1. We previously reported transcriptomic and proteomic evidence for differential expression (DE) patterns between segregating F1 hybrid avirulent and virulent aphid populations and between their parental genotypes (Thorpe et al., 2024). In that study, CHL1 was the only gene/protein found to be DE across multiple experiments including RNA-Seq of heads and whole bodies, and saliva proteomics of parental genotypes, in all cases being upregulated in avirulent aphids. Here we established that CHL1 protein abundance co-segregates with avirulence through comparative proteomics of saliva of multiple representatives of the virulent and avirulent sides of the F1 hybrid populations. In all cases, CHL1 protein was present in saliva of avirulent individuals but undetectable in virulent counterparts. RNA-Seq analysis revealed alternative splicing of CHL1, with the long isoform expressed only in avirulent genotypes. Moreover, an exon SNP within the long isoform enabled deduction that the long isoform was expressed only from one of the alleles originating from the avirulent parent clone PS01. This SNP was absent from the virulent parent N116 and from virulent F1 progeny. Although considerable time and effort are required to create and analyse aphid F1 populations, the controlled genetic approach provides a powerful means to associate effector expression with aphid (a)virulence phenotypes on a host carrying a known resistance QTL, *RAP1* (Stewart et al., 2009). In contrast, many more DE genes are found between the divergent parents, that are adapted to different legume host species, in this case *Pisum sativum* (garden pea) and *Medicago sativa* (alfalfa).,Most of the differentials between these parents may thus be attributable to aphid genomic background rather than causatively related to avirulence.

The short isoform of CHL1 appears to be a loss of function mutation, resulting in inability to express or secrete CHL1 protein. The absence of the CHL1 long isoform in the virulent parent clone N116, a clone that falls within the biotype class that has a preference for alfalfa hosts, could represent a component of host adaptation if such hosts have a cognate receptor that detects CHL1 protein. Lack of expression of functional CHL1 would not necessarily impair aphid fitness if the other CHL family proteins provide redundancy. The causative R gene underlying the *RAP1* QTL is not yet defined but may be an NLR immune receptor. Because CHL1 acts as an avirulence factor on hosts carrying *RAP1,* the outcome appears to conform to the gene-for-gene model, resulting in an ETI response. It remains to be discovered whether CHL1 interacts directly with *RAP1* gene products or indirectly via other host proteins.

### Conserved gene and protein structure in the CHL family

Analysis of exon-intron patterns, protein sequences and predicted protein structures indicated strong conservation of several features across the CHL family. Most genes are comprised of eight exons, encoding proteins of 342 to 371 amino acids in *A. pisum*. Even for the most distant relatives that carry CHLs, such as *Daktulosphaira vitifolia*, the single gene in that species has the canonical eight exon structure and a predicted protein of 359 amino acids. It can be inferred that CHL gene organisation has most likely retained an ancestral state, dating back to the separation of aphids from sister groups The AlphaFold models were also highly similar across all CHLs in *A. pisum* (Fig. 3E), with high confidence mean Template Modelling score of 0.861 and RMSD 1.76 Å, despite the low mean sequence identity (22%). Some CHLs are expressed as truncated forms, either from tandem duplicate genes or alternatively spliced isoforms, resulting in protein lengths of 258 to 306 amino acids. We speculate that the shorter proteins may lack biochemical functions, but this remains to be experimentally determined. For *A. pisum* at least, all CHLs found in saliva contain predicted secretion signals, either a classic signal peptide (SP) or an unconventional (UPS) type (Table 3). The short isoform of ApCHL1 lacks SP and UPS, and although transcribed, was not found as protein in saliva.

### Origin and expansion of the CHL family

Based on the presence of CHL proteins only in the genomes of Aphidomorpha, we can infer that the CHL family originated subsequent to divergence from ancient sister groups. This separation occurred over 200 million years ago (Julca et al., 2020), hence CHLs can be considered an ancient protein family. This time also overlaps with the estimated dates for emergence of the angiosperms (Ramírez-Barahona et al., 2020; Zuntini et al., 2024). Prior to this event, fossil evidence indicates aphid groups fed on other vascular plants including gymnosperms (Labandeira et al., 2016; Labandeira and Wappler, 2023). Because CHLs are found in extant gymnosperm-feeding genera such as *Cinara* and *Adelges*, it is unlikely that CHLs are necessary only for species living on angiosperms.

Although CHL biochemical functions have yet to be experimentally determined, some ideas are explored here. First, CHL structural models predict the potential to bind chitin oligomer ligands (Fig. 5B) within the major cleft of CHLs, and with an orientation somewhat similar to the known positioning of chitin binding of true chitinases (Fig. 5C-E). However, CHLs all lack the DxxDxDxE conserved catalytic motif of chitinases, so although they may bind chitin, it is possible that CHLs are not enzymatically active. There are defined other chitin-binding domains (CBDs; InterPro IPR002557) in chitinases and some other proteins of invertebrates and plants (Suetake et al., 2000), but these were not found for CHL proteins. A second point to note is that there are other groups of phloem-feeding within the Hemiptera, including psyllids and mealybugs, that do not possess the CHL protein family. Therefore, CHLs are not necessarily fundamental to aphids’ ability to successfully locate and feed from phloem.

Psyllids such as *D. citri*, the vector of the causative organism for the important citrus greening disease (Hall et al., 2013), have proteins also described as chitinase-like (Wu et al., 2022). However, these chitinase-like proteins sit within the true chitinase and IDGF clades and appear unrelated to aphid CHLs. Indeed, expression of only one *D. citri* chitinase-like gene was found to be upregulated in salivary glands, and the functions were proposed instead to relate to insect development (Wu et al., 2022).

Numbers of CHLs detected in species with high quality genome data vary considerably. The sister groups to true aphids, Phylloxeroidea and Adelgoidea, possess one or two, respectively. Based on searches across well-curated genomes within the Aphidoidea, gene numbers range from two in *D. noxia* to more than 40 in *R. padi*. The evolutionary drivers for CHL family expansion are presently unclear, but increased gene numbers are likely to provide redundancy, assuming that CHLs share the same or similar biochemical functions when acting as virulence factors.

### Aphid CHL proteins may have analogous functions to fungal chitinase-like effectors

We hypothesise above that aphid CHL proteins may retain chitin-binding properties. On this basis, CHLs could act to sequester chitin and chitin oligosaccharide fragments, suppressing the classic PTI immune response that acts through cell surface chitin receptors such as MtCERK1/LYK9 and MtLYR4 (Feng et al., 2019). Alternatively, CHL proteins could out-compete secreted host chitinases that otherwise may attack the aphid’s chitin structures. Both these virulence functions would likely occur in the extracellular space. Intriguingly, there are striking parallels for both such modes of action in chitin-binding fungal effectors: for example, *Cladosporium fulvum* CfEcp6 suppresses chitin perception by chitin PAMP receptors (Sánchez-Vallet et al., 2013) whereas CfAvr4 blocks chitinases from accessing their substrates (van den Burg et al., 2006). These fungal effectors contain lysM chitin binding domains, notably absent from CHLs. However, another chitinase-like fungal effector from *Moniliophthora perniciosa* (MpChi) (Fiorin et al., 2018) is related to fungal chitinases but not to aphid chitinases or to aphid CHLs. It remains to be established whether aphid CHLs and these fungal effectors are examples of functional evolutionary convergence.

Despite the wealth of evidence for immune activation following detection of fungal chitin, surprisingly there is scant evidence to support a role for chitin in detection of invertebrate attack. Notably, Arabidopsis *CERK1* receptor mutants were not compromised in defence responses to aphid-derived extracts, although other unknown PRRs and the BAK1 co-receptor were implicated (Prince et al., 2014). However, one recent paper points to chitin oligomers in oral secretions of chewing herbivores being responsible for triggering CERK1-dependent defences in rice (Kanda et al., 2025).

## Conclusion

From the extensive list of uncharacterised candidate effectors present in aphid saliva, we have uncovered a hitherto unrecognised family of chitinase-like proteins, present only in aphids. In pea aphid, several CHLs are amongst the most abundant salivary proteins, likely indicating functional importance. We propose that CHLs evolved as virulence factors, possibly acting to suppress chitin-triggered immunity. The presence of multiple CHLs in saliva further indicates likely redundancy. Moreover, the founding member, ApCHL1, additionally appears to have an avirulence function, associated with aphid incompatibility on host plants carrying the *RAP1* resistance locus. We conclude that loss of ApCHL1 protein expression in our virulent genotypes does not appear to impair fitness on hosts lacking *RAP1.* Future studies can explore the biochemical basis of CHL functions. For CHL virulence, testing affinity for chitin would be a logical first step. For the specific avirulence function of ApCHL1, detection by a host protein is predicted, either directly by *RAP1* or indirectly by another associated receptor or host target.

### Methods Materials

Pea aphid (*Acyrthosiphon pisum*) clones were maintained on tic bean (*Vicia faba minor*) or broad bean (*Vicia faba*) cv. The Sutton plants as previously described (Kanvil et al., 2015; Thorpe et al., 2024). Host plants for experiments were tic bean or *Medicago truncatula* genotypes Jemalong A17 and DZA315.16.

### Gene expression analysis

Gene expression was measured by qRT-PCR. Target gene expression was calculated relative to the average expression of two reference genes: GAPDH (glyceraldehyde-3-phosphate dehydrogenase; ACYPI009769) and SDHB (succinate dehydrogenase B; ACYPI003519). Primer sequences are listed in Supplementary Table S12. Expression values were calculated by the Livak ΔΔCt method. For all experiments, three independent biological replicates were analysed, with three technical replicates.

### Proteomics

### a. Label-free quantitative proteomics

Diet collected saliva samples were obtained from the VIR (N116) and AVR (PS01) parental clones in addition to their F1 VIR (PN35) and AVR (PN1) offspring that were established from single individuals selected from the F1 populations reported previously (Thorpe et al., 2024). Saliva collection was done essentially as described previously (Thorpe et al., 2024). Briefly, aphids were transferred to Perspex rings (radius 4.5 cm, height 5 cm), each containing 4.5 ml of chemically defined diet held between two stretched sheets of Parafilm™. The aphids were reared on the diets at 20°C for 24 h after which the diets were pooled and frozen at -80°C. Samples were concentrated using a Vivacell 250 Pressure Concentrator (Sartorius Mechatronics, UK) using a 5000 Da molecular weight cut-off (MWCO) polyethersulfone (PES) membrane, then Roche cOmplete™ protease inhibitor cocktail (PIC) was added. After further concentration to approximately 250 μl using a Vivaspin 6 centrifuge concentrator (Sartorius Mechatronics, UK) with a 5000 Da MWCO PES membrane, samples were purified using a 2D Clean-up Kit (GE HealthCare) then protein pellets were re suspended in 25 μl 6 M urea, 2 M thiourea, 0.1 M Tris–HCl, pH 8.0. After reduction with dithiothreitol and alkylation with iodoacetamide, proteins were digested with Sequence Grade Trypsin (Promega) and ProteaseMax Surfactant Trypsin Enhancer (Promega) at 37°C for 18 h.

Mass spectrometry and data analysis were done as previously described (Thorpe et al., 2024). Three independent biological replicates per genotype were analysed. Briefly, digested peptides were loaded onto a Dionex Ultimate 3000 (RSLCnano) chromatography system connected to a QExactive (ThermoFisher Scientific) mass spectrometer. Peptides were separated by a 50 min acetonitrile gradient on a Biobasic C18 PicofritTM column (100 mm length, 75 µm ID), at flow rate 250 nl min^−1^. Data were acquired in automatic data dependent switching mode. A high-resolution MS scan (300– 2000 Da) was performed using the Orbitrap to select the 15 most intense ions prior to MS/MS.

Protein identification was done with MaxQuant (version 2.4.2.0; http://maxquant.org/) to correlate the data against the previously reported predicted protein set (ACPISUM_Proteins; 30,891 entries) (Thorpe et al., 2024) using default search parameters for Orbitrap data. False Discovery Rates were set to 1% for both peptides and proteins and the FDR was estimated following searches against a target-decoy database. Quantitative analyses were conducted in Perseus (Version 2.0.10.0 http://maxquant.org/) using the normalized label-free quantitation (LFQ) intensity values.

The data were filtered to remove contaminants, reverse proteins (identified from peptides derived from the reversed part of the decoy database) and peptides identified by site. LFQ intensity values were log2 transformed and samples were allocated to their corresponding groups to determine the qualitative distribution of proteins of interest. Proteins not found in all three replicates of at least one sample were omitted from the analysis.

### b. LCMS-Multiple Reaction Monitoring (MRM) protein quantitation

Three representative VIR (NP10, PN14, PN35) and AVR (NP18, NP22, PN20) clones were selected from the F1 populations reported previously (Thorpe et al., 2024), along with the parental genotypes N116 and PS01. Three independent saliva samples were collected for two days from each clone via sterilised aphid diets (2 ml of 15% (w/v) sucrose; 100 mM L-serine; 100 mM L-methionine; 100 mM L-aspartic acid, pH 7.2), similar to description above. Approximately 4000 aphids were cultured for each saliva replicate, with around 200 aphids aged four to seven days per dish. Green translucent plastic sheets were suspended over the petri dishes to mimic leaves and encourage the aphids to feed. Proteins were concentrated with VivaspinⓇ 15R Centrifugal Concentrators, 5000 MWCO HydrosartⓇ (Sartorius, Germany) solution. Concentrators were first flushed with 5 to 10 ml of sterile PBS (137 mM NaCl; 10 mM Phosphate; 2.7 mM KCl, pH 7.4), then 10 ml salivary solution was added, followed by a repeat wash with 5 to 10 ml PBS. Approximately 200 to 300 μl of the resultant concentrated supernatant was further reduced in volume with AmiconⓇ Ultra-0.5 Centrifugal Filters, 3000 NMWL (Merck, Germany). Protein concentrations were quantified at 280 nm using Nanodrop (ThermoFisher, USA).

A scheduled MRM method was developed for detection of unique peptides from ApCHL1 protein. Peptides for spectrum acquisition were selected by *in silico* trypsin digestion and information-dependent acquisition (IDA) LC-MS experiments. The MRM detection window for each candidate peptide was centred on the retention time for each ion fragment, enabling the higher sensitivity of a scheduled MRM method. For method development, the full length ApCHL1 coding sequence was cloned into destination vector Gateway® Nova pET-53-DEST™ (Merck, Germany) by Gateway™ LR reaction. Recombinant ApCHL1 protein was expressed in *E. coli* BL21 cells at 37 °C for four to six hours, following transformation and induction with IPTG (1 mM). Total protein was extracted in 6 M urea; 2 M thiourea; 0.2 M Tris-HCl, pH 8.0. Recombinant proteins from *E. coli* extracts (30 μl) were reduced in 3 μl of 100 mM Tris(2-carboxyethyl)phosphine hydrochloride (TCEP) and 260 μl of 50 mM NH4CO3 buffer. Samples were digested by Trypsin-ultra™, Mass Spectrometry Grade (NEB, USA) overnight at 37°C and the reaction stopped by addition of 15 μl 5% formic acid the following day. Samples submitted for LC-MS analysis contained approximately 1 M urea, 1 mM TCEP and 40 mM NH4CO3

Protein concentrations of saliva samples were adjusted to 20 μg in 75 μl of 50mM NH4CO3 of resuspension buffer (6 M urea; 2 M thiourea; 0.1 M Tris-HCl, pH 8.0). In-solution digestion was performed by first suspending samples in 105 μl 50 mM NH4CO3, reducing samples in 0.55 M DTT for 30 minutes at 56°C and alkylating with 10.5 M IAA for 20 minutes at room temperature. Next, 1 μl of 1% ProteaseMAX ™ Surfactant stock (Promega, USA) and 1 μl of 0.5 μg μl^-1^ Trypsin-ultra™, Mass Spectrometry Grade (NEB, USA) were added and samples incubated overnight at 37°C. The reaction was stopped with 1 μl TFA at room temperature. Protein mixtures were centrifuged at 13,000 *g* and dried using an Eppendorf Concentrator Plus (Eppendorf, UK) at 30°C for approximately two to three hours. Pellets were resuspended in 100 μl LC-MS buffer (0.1% TFA; 2% acetonitrile in HPLC water), with final protein concentrations of approximately 0.2 μg μl^-1^.

Samples were analysed by LC-MS using a Shimadzu LC20/CBM20A binary LC system (Shimadzu) and a QTRAPⓇ 6500 MS [AB Sciex]) in scheduled MRM positive ion mode. Column was Luna C18 3 mm x 100 mm Phenomenex Luna 3 µm C18(2) 100 mm x 2 mm, with flow rate of 250 µl min^-1^. Mobile phase comprised 94.9% H2O, 5% CH3CN, 0.1% CHOOH (solvent A) and 94.9% CH3CN, 5% H2O, 0.1% CHOOH (solvent B). Initial conditions were 100% A, followed by a linear gradient to 25% B over 30 min, then to 50% B at 35 min and 100% B at 37 min. Data analysis was conducted with Analyst software (version 1.6.3) by building a quantitation method for ApCHL1, based on the strongest peptide signal (YIDTIDPTAK.+2y8; Q1 543.811, Q3 716.430 m/z) with the second strongest peptide (EIQAQGIVTK.+2y7; Q1 543.811, Q3 716.430 m/z) used for confirmation. Instrument performance was checked with a peptide fragment ion IGGIGTVPVGR.+2y7 (Q1 513.309, Q3 685.399 m/z) from reference protein elongation factor 1-alpha (LOC100574365)

### Bioinformatics

Homologs of ACPISUM_029930 were initially detected by BLASTP searches at NCBI using default settings, and an E value cutoff of 10^-5^. Weaker homologs were detected by PSI-BLAST at NCBI using three to five cycles, with either default settings or with E value cutoff set at 1 or 0.1.

Neighbour-joining unrooted trees were created using the tree construction module within NCBI Blast suite. Manual curation of annotated isoforms retained only the longest form, and short predicted proteins (<250 amino acids) were also removed. Dissimilarity cutoff was default 0.8 for BLASTP results and set to 0.9 for PSI-BLAST.

WebLogo (Crooks et al., 2004) was used to highlight conserved residues, and conserved regions were identified from COBALT alignments.

AlphaFold 2 and AlphaFold 3 (Abramson et al., 2024; Jumper et al., 2021) were used to generate predicted protein models, which were visualised in PyMol (DeLano et al., n.d.; Schrödinger, LLC, 2015). RAPTOR-X Deep Align was used to align multiple proteins, with confidence of outputs being based on TM score and RMSD (Å) (Wang et al., 2013). Homologous structures were identified from searches via Dali and FoldSeek.

For molecular docking studies, the SwissDock web server was used (Grosdidier et al., 2011). Tetrameric chitin was used as the query ligand, the structure of which was obtained from PubChem at https://pubchem.ncbi.nlm.nih.gov/compound/5288898.

### Data availability

The mass spectrometry proteomics data have been deposited to the ProteomeXchange Consortium via the PRIDE (Perez-Riverol et al., 2025) partner repository with the dataset identifier PXD066490.

## Acknowledgements

We thank the Biotechnology and Biological Sciences Research Council for funding to CT (BB/N002830/1 AND BB/X002322/1) and JB (BB/N002660/1). We acknowledge expert technical support from Mark H. Bennett, Paul Hitchen and Ji-Won Min.

## Author contributions

CT, JB and JC conceived the study and designed experiments. RLC, SA, PT, ND, PYS, LE, ML, EH, EK and JI designed and conducted experiments. RLC, SA, PT ND, PYS, LE, ML, EH, EK, JI, CT and JC analysed data. CT, JB and JC wrote the paper. All authors approved the final manuscript.

## Notes

### Competing Interest Statement

The authors have declared no competing interest.

## References

Abramson, J., et al. 2024. Accurate structure prediction of biomolecular interactions with AlphaFold 3. Nature 630, 493–500. 10.1038/s41586-024-07487-w

Bos, J.I.B., Prince, D., Pitino, M., Maffei, M.E., Win, J., Hogenhout, S.A., 2010. A functional genomics approach identifies candidate effectors from the aphid species *Myzus persicae* (Green Peach Aphid). PLOS Genet. 6, e1001216. 10.1371/journal.pgen.1001216

Carolan, J.C., Caragea, D., Reardon, K.T., Mutti, N.S., Dittmer, N., Pappan, K., Cui, F., Castaneto, M., Poulain, J., Dossat, C., Tagu, D., Reese, J.C., Reeck, G.R., Wilkinson, T.L., Edwards, O.R., 2011. Predicted effector molecules in the salivary secretome of the pea aphid (*Acyrthosiphon pisum*): A dual transcriptomic/proteomic approach. J. Proteome Res. 10, 1505–1518. 10.1021/pr100881q

Carolan, J.C., Fitzroy, C.I.J., Ashton, P.D., Douglas, A.E., Wilkinson, T.L., 2009. The secreted salivary proteome of the pea aphid *Acyrthosiphon pisum* characterised by mass spectrometry. Proteomics 9, 2457–2467. 10.1002/pmic.200800692

Chaudhary, R., Atamian, H.S., Shen, Z., Briggs, S.P., Kaloshian, I., 2015. Potato aphid salivary proteome: enhanced salivation using resorcinol and identification of aphid phosphoproteins. J. Proteome Res. 14, 1762–1778. 10.1021/pr501128k

Chaudhary, R., Atamian, H.S., Shen, Z., Briggs, S.P., Kaloshian, I., 2014. GroEL from the endosymbiont *Buchnera aphidicola* betrays the aphid by triggering plant defense. Proc. Natl. Acad. Sci. U. S. A. 111, 8919–8924. 10.1073/pnas.1407687111

Chaudhary, R., Peng, H.-C., He, J., MacWilliams, J., Teixeira, M., Tsuchiya, T., Chesnais, Q., Mudgett, M.B., Kaloshian, I., 2019. Aphid effector Me10 interacts with tomato TFT7, a 14-3-3 isoform involved in aphid resistance. New Phytol. 221, 1518–1528. 10.1111/nph.15475

Crooks, G.E., Hon, G., Chandonia, J.-M., Brenner, S.E., 2004. WebLogo: A sequence logo generator. Genome Res. 14, 1188–1190. 10.1101/gr.849004

Dogimont, C., Chovelon, V., Pauquet, J., Boualem, A., Bendahmane, A., 2014. The *Vat* locus encodes for a CC-NBS-LRR protein that confers resistance to *Aphis gossypii* infestation and *A. gossypii-* mediated virus resistance. Plant J. 80, 993–1004. 10.1111/tpj.12690

Du, B., Zhang, W., Liu, B., Hu, J., Wei, Z., Shi, Z., He, R., Zhu, L., Chen, R., Han, B., He, G., 2009. Identification and characterization of *Bph14*, a gene conferring resistance to brown planthopper in rice. Proc. Natl. Acad. Sci. 106, 22163–22168. 10.1073/pnas.0912139106

Elzinga, D.A., De Vos, M., Jander, G., 2014. Suppression of plant defenses by a *Myzus persicae* (Green Peach Aphid) salivary effector protein. Mol. Plant-Microbe Interactions® 27, 747–756. 10.1094/MPMI-01-14-0018-R

Feng, F., Sun, J., Radhakrishnan, G.V., Lee, T., Bozsóki, Z., Fort, S., Gavrin, A., Gysel, K., Thygesen, M.B., Andersen, K.R., Radutoiu, S., Stougaard, J., Oldroyd, G.E.D., 2019. A combination of chitooligosaccharide and lipochitooligosaccharide recognition promotes arbuscular mycorrhizal associations in *Medicago truncatula*. Nat. Commun. 10, 5047. 10.1038/s41467-019-12999-5

Fiorin, G.L., Sanchéz-Vallet, A., Thomazella, D.P. de T., do Prado, P.F.V., do Nascimento, L.C., Figueira, A.V. de O., Thomma, B.P.H.J., Pereira, G.A.G., Teixeira, P.J.P.L., 2018. Suppression of plant immunity by fungal chitinase-like effectors. Curr. Biol. 28, 3023–3030.e5. 10.1016/j.cub.2018.07.055

Gravino, M., Mugford, S.T., Kreuter, N., Joyce, J., Wilson, C., Kliot, A., Canham, J., Mathers, T.C., Drurey, C., Maqbool, A., Martins, C., Saalbach, G., Hogenhout, S.A., 2024. The conserved aphid saliva chemosensory protein effector Mp10 targets plant AMSH deubiquitinases at cellular membranes to suppress pattern-triggered immunity. 10.1101/2024.11.15.622802

Grosdidier, A., Zoete, V., Michielin, O., 2011. SwissDock, a protein-small molecule docking web service based on EADock DSS. Nucleic Acids Res. 39, W270–W277. 10.1093/nar/gkr366

Guo, J., Wang, H., Guan, W., Guo, Q., Wang, J., Yang, J., Peng, Y., Shan, J., Gao, M., Shi, S., Shangguan, X., Liu, B., Jing, S., Zhang, J., Xu, C., Huang, J., Rao, W., Zheng, X., Wu, D., Zhou, C., Du, B., Chen, R., Zhu, L., Zhu, Y., Walling, L.L., Zhang, Q., He, G., 2023. A tripartite rheostat controls self-regulated host plant resistance to insects. Nature 618, 799–807. 10.1038/s41586-023-06197-z

Hall, D.G., Richardson, M.L., Ammar, E.-D., Halbert, S.E., 2013. Asian citrus psyllid, *Diaphorina citri*, vector of citrus huanglongbing disease. Entomol. Exp. Appl. 146, 207–223. 10.1111/eea.12025

Julca, I., Marcet-Houben, M., Cruz, F., Vargas-Chavez, C., Johnston, J.S., Gómez-Garrido, J., Frias, L., Corvelo, A., Loska, D., Cámara, F., Gut, M., Alioto, T., Latorre, A., Gabaldón, T., 2020. Phylogenomics identifies an ancestral burst of gene duplications predating the diversification of Aphidomorpha. Mol. Biol. Evol. 37, 730–756. 10.1093/molbev/msz261

Jumper, J., et al. 2021. Highly accurate protein structure prediction with AlphaFold. Nature 596, 583–589. 10.1038/s41586-021-03819-2

Kanda, Y., Shinya, T., Wari, D., Hojo, Y., Fujiwara, Y., Tsuchiya, W., Fujimoto, Z., Thomma, B.P.H.J., Nishizawa, Y., Kamakura, T., Galis, I., Mori, M., 2025. Chitin-signaling-dependent responses to insect oral secretions in rice cells propose the involvement of chitooligosaccharides in plant defense against herbivores. Plant J. 121, e17157. 10.1111/tpj.17157

Kanvil, S., Collins, C.M., Powell, G., Turnbull, C.G.N., 2015. Cryptic virulence and avirulence alleles revealed by controlled sexual recombination in pea aphids. Genetics 199, 581–593. 10.1534/genetics.114.173088

Labandeira, C.C., Kustatscher, E., Wappler, T., 2016. Floral assemblages and patterns of insect herbivory during the Permian to Triassic of Northeastern Italy. PLOS ONE 11, e0165205. 10.1371/journal.pone.0165205

Labandeira, C.C., Wappler, T., 2023. Arthropod and pathogen damage on fossil and modern plants: exploring the origins and evolution of herbivory on land. Annu. Rev. Entomol. 68, 341–361. 10.1146/annurev-ento-120120-102849

Pavithran, S., Murugan, M., Yogendra, K., Mannu, J., Venkatasamy, B., Sanivarapu, H., Harish, S., Natesan, S., Onkarappa, D., 2024. Proteomic insights into the saliva and salivary glands of the cotton aphid, *Aphis gossypii* (Hemiptera: Aphididae). Phytoparasitica 52, 73. 10.1007/s12600-024-01192-0

Peng, H.-C., Mantelin, S., Hicks, G.R., Takken, F.L.W., Kaloshian, I., 2016. The conformation of a plasma membrane-localized somatic embryogenesis receptor kinase complex is altered by a potato aphid-derived effector. Plant Physiol. 171, 2211–2222. 10.1104/pp.16.00295

Perez-Riverol, Y., Bandla, C., Kundu, D.J., Kamatchinathan, S., Bai, J., Hewapathirana, S., John, N.S., Prakash, A., Walzer, M., Wang, S., Vizcaíno, J.A. The PRIDE database at 20 years: 2025 update. Nucleic Acids Res. 2025 53, D543–D553. 10.1093/nar/gkae1011.

Pitino, M., Hogenhout, S.A., 2013. Aphid protein effectors promote aphid colonization in a plant species-specific manner. Mol. Plant-Microbe Interactions® 26, 130–139. 10.1094/MPMI-07-12-0172-FI

Prince, D.C., Drurey, C., Zipfel, C., Hogenhout, S.A., 2014. The leucine-rich repeat receptor-like kinase BRASSINOSTEROID INSENSITIVE1-ASSOCIATED KINASE1 and the cytochrome P450 PHYTOALEXIN DEFICIENT3 contribute to innate immunity to aphids in Arabidopsis. Plant Physiol. 164, 2207–2219. 10.1104/pp.114.235598

Ramírez-Barahona, S., Sauquet, H., Magallón, S., 2020. The delayed and geographically heterogeneous diversification of flowering plant families. Nat. Ecol. Evol. 4, 1232–1238. 10.1038/s41559-020-1241-3

Rao, W., Ma, T., Cao, J., Zhang, Y., Chen, S., Lin, S., Liu, X., He, G., Wan, L., 2024. Recognition of a salivary effector by the TNL protein RCSP promotes effector-triggered immunity and systemic resistance in *Nicotiana benthamiana*. J. Integr. Plant Biol. n/a. 10.1111/jipb.13800

Rodriguez, P.A., Escudero-Martinez, C., Bos, J.I.B., 2017. An aphid effector targets trafficking protein VPS52 in a host-specific manner to promote virulence. Plant Physiol. 173, 1892–1903. 10.1104/pp.16.01458

Rossi, M., Goggin, F.L., Milligan, S.B., Kaloshian, I., Ullman, D.E., Williamson, V.M., 1998. The nematode resistance gene *Mi* of tomato confers resistance against the potato aphid. Proc. Natl. Acad. Sci. 95, 9750–9754. 10.1073/pnas.95.17.9750

Sánchez-Vallet, A., Saleem-Batcha, R., Kombrink, A., Hansen, G., Valkenburg, D.-J., Thomma, B.P., Mesters, J.R., 2013. Fungal effector Ecp6 outcompetes host immune receptor for chitin binding through intrachain LysM dimerization. eLife 2, e00790. 10.7554/eLife.00790

Schrödinger, LLC, 2015. The PyMOL Molecular Graphics System, Version 1.8.

Schultz-Larsen, T., Lenk, A., Kalinowska, K., Vestergaard, L.K., Pedersen, C., Isono, E., Thordal-Christensen, H., 2018. The AMSH3 ESCRT-III-associated deubiquitinase is essential for plant immunity. Cell Rep. 25, 2329–2338.e5. 10.1016/j.celrep.2018.11.011

Stewart, S.A., Hodge, S., Ismail, N., Mansfield, J.W., Feys, B.J., Prospéri, J.-M., Huguet, T., Ben, C., Gentzbittel, L., Powell, G., 2009. The *RAP1* gene confers effective, race-specific resistance to the pea aphid in *Medicago truncatula* independent of the hypersensitive reaction. Mol. Plant-Microbe Interactions® 22, 1645–1655. 10.1094/MPMI-22-12-1645

Suetake, T., Tsuda, S., Kawabata, S., Miura, K., Iwanaga, S., Hikichi, K., Nitta, K., Kawano, K., 2000. Chitin-binding proteins in invertebrates and plants comprise a common chitin-binding structural Motif. J. Biol. Chem. 275, 17929–17932. 10.1074/jbc.C000184200

Thorpe, P., Altmann, S., Lopez-Cobollo, R., Douglas, N., Iqbal, J., Kanvil, S., Simon, J.-C., Carolan, J.C., Bos, J., Turnbull, C., 2024. Multi-omics approaches define novel aphid effector candidates associated with virulence and avirulence phenotypes. BMC Genomics 25, 1065. 10.1186/s12864-024-10984-x

Thorpe, P., Cock, P.J.A., Bos, J., 2016. Comparative transcriptomics and proteomics of three different aphid species identifies core and diverse effector sets. BMC Genomics 17, 172. 10.1186/s12864-016-2496-6

van Bel, A.J.E., Will, T., 2016. Functional evaluation of proteins in watery and gel saliva of aphids. Front. Plant Sci. 7. 10.3389/fpls.2016.01840

van den Burg, H.A., Harrison, S.J., Joosten, M.H.A.J., Vervoort, J., de Wit, P.J.G.M., 2006. *Cladosporium fulvum* Avr4 protects fungal cell walls against hydrolysis by plant chitinases accumulating during infection. Mol. Plant-Microbe Interactions® 19, 1420–1430. 10.1094/MPMI-19-1420

Waksman, T., Astin, E., Fisher, S.R., Hunter, W.N., Bos, J.I.B., 2024. Computational prediction of structure, function, and interaction of *Myzus persicae* (Green Peach Aphid) salivary effector proteins. Mol. Plant-Microbe Interactions® 37, 338–346. 10.1094/MPMI-10-23-0154-FI

Wang, S., Ma, J., Peng, J., Xu, J., 2013. Protein structure alignment beyond spatial proximity. Sci. Rep. 3, 1448. 10.1038/srep01448

Wu, C.-H., Abd-El-Haliem, A., Bozkurt, T.O., Belhaj, K., Terauchi, R., Vossen, J.H., Kamoun, S., 2017. NLR network mediates immunity to diverse plant pathogens. Proc. Natl. Acad. Sci. 114, 8113–8118. 10.1073/pnas.1702041114

Wu, Z.-Z., Zhang, W.-Y., Lin, Y.-Z., Li, D.-Q., Shu, B.-S., Lin, J.-T., 2022. Genome-wide identification, characterization and functional analysis of the chitinase and chitinase-like gene family in *Diaphorina citri*. Pest Manag. Sci. 78, 1740–1748. 10.1002/ps.6793

Yu, X.-D., Liu, Z.-C., Huang, S.-L., Chen, Z.-Q., Sun, Y.-W., Duan, P.-F., Ma, Y.-Z., Xia, L.-Q., 2016. RNAi-mediated plant protection against aphids. Pest Manag. Sci. 72, 1090–1098. 10.1002/ps.4258

Zhang, H., Lin, R., Liu, Q., Lu, J., Qiao, G., Huang, X., 2023. Transcriptomic and proteomic analyses provide insights into host adaptation of a bamboo-feeding aphid. Front. Plant Sci. 13. 10.3389/fpls.2022.1098751

Zhang, Y., Fu, Y., Francis, F., Liu, X., Chen, J., 2021. Insight into watery saliva proteomes of the grain aphid, *Sitobion avenae*. Arch. Insect Biochem. Physiol. 106, e21752. 10.1002/arch.21752

Zuntini, A.R., et al. 2024. Phylogenomics and the rise of the angiosperms. Nature 629, 843–850. 10.1038/s41586-024-07324-0

