## Supplemental Figures S1 to S8 for "A novel chitinase-like family of candidate effectors unique to aphids"

### Slide 1
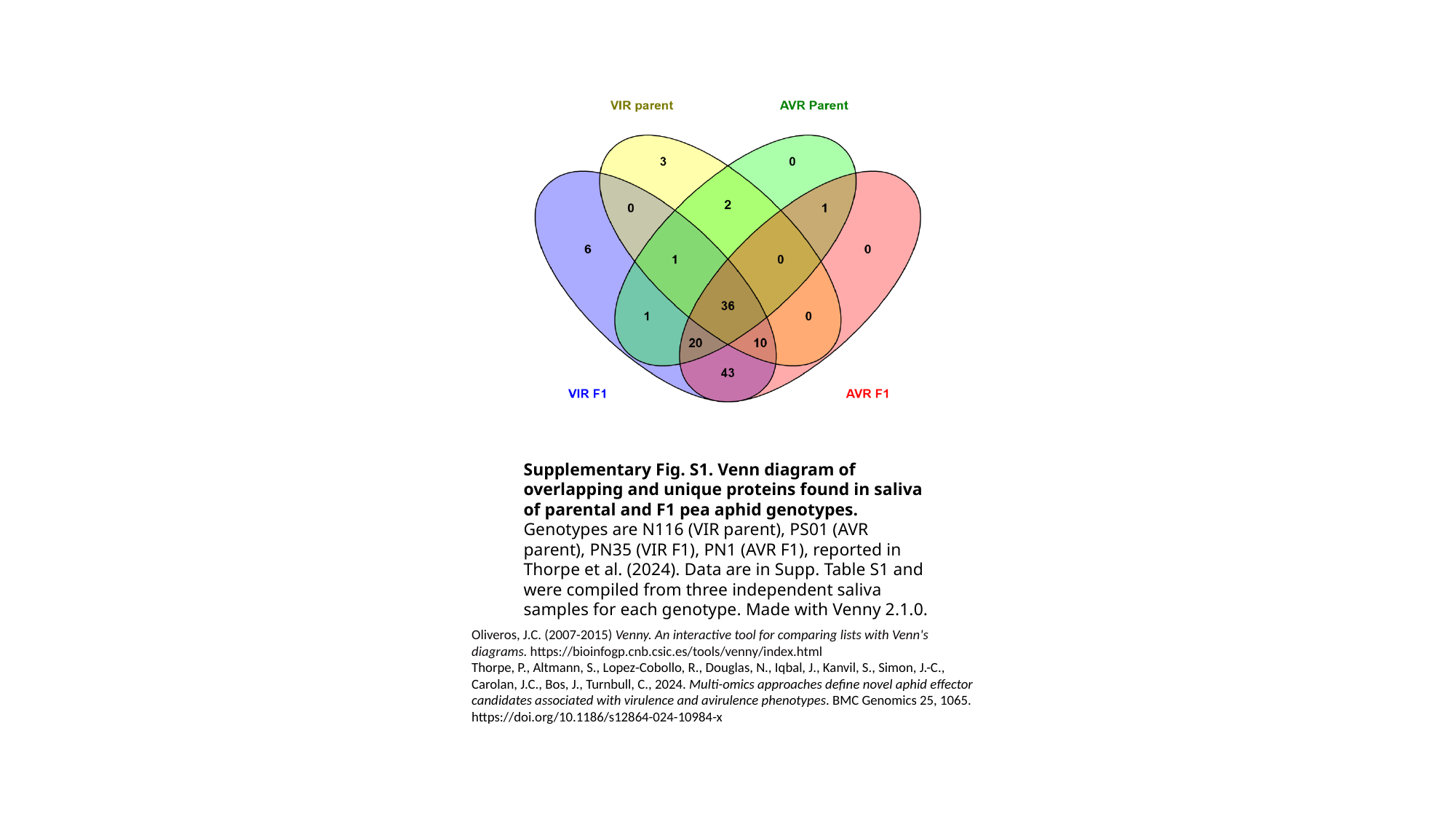

Supplementary Fig. S1. Venn diagram of overlapping and unique proteins found in saliva of parental and F1 pea aphid genotypes. Genotypes are N116 (VIR parent), PS01 (AVR parent), PN35 (VIR F1), PN1 (AVR F1), reported in Thorpe et al. (2024). Data are in Supp. Table S1 and were compiled from three independent saliva samples for each genotype. Made with Venny 2.1.0.
Oliveros, J.C. (2007-2015) Venny. An interactive tool for comparing lists with Venn's diagrams. https://bioinfogp.cnb.csic.es/tools/venny/index.html
Thorpe, P., Altmann, S., Lopez-Cobollo, R., Douglas, N., Iqbal, J., Kanvil, S., Simon, J.-C., Carolan, J.C., Bos, J., Turnbull, C., 2024. Multi-omics approaches define novel aphid effector candidates associated with virulence and avirulence phenotypes. BMC Genomics 25, 1065. https://doi.org/10.1186/s12864-024-10984-x

### Slide 2
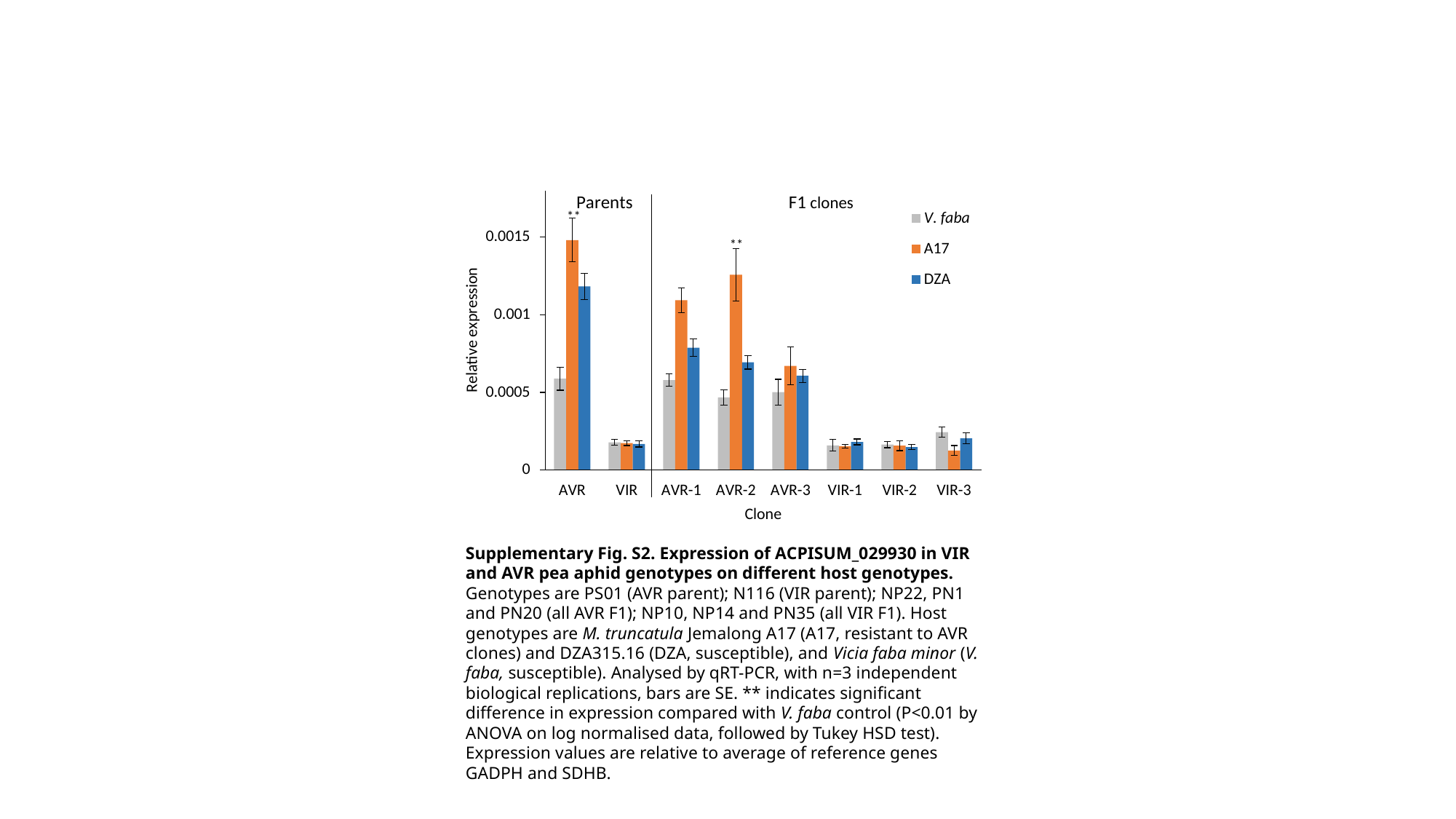

Supplementary Fig. S2. Expression of ACPISUM_029930 in VIR and AVR pea aphid genotypes on different host genotypes. Genotypes are PS01 (AVR parent); N116 (VIR parent); NP22, PN1 and PN20 (all AVR F1); NP10, NP14 and PN35 (all VIR F1). Host genotypes are M. truncatula Jemalong A17 (A17, resistant to AVR clones) and DZA315.16 (DZA, susceptible), and Vicia faba minor (V. faba, susceptible). Analysed by qRT-PCR, with n=3 independent biological replications, bars are SE. ** indicates significant difference in expression compared with V. faba control (P<0.01 by ANOVA on log normalised data, followed by Tukey HSD test). Expression values are relative to average of reference genes GADPH and SDHB.

### Slide 3
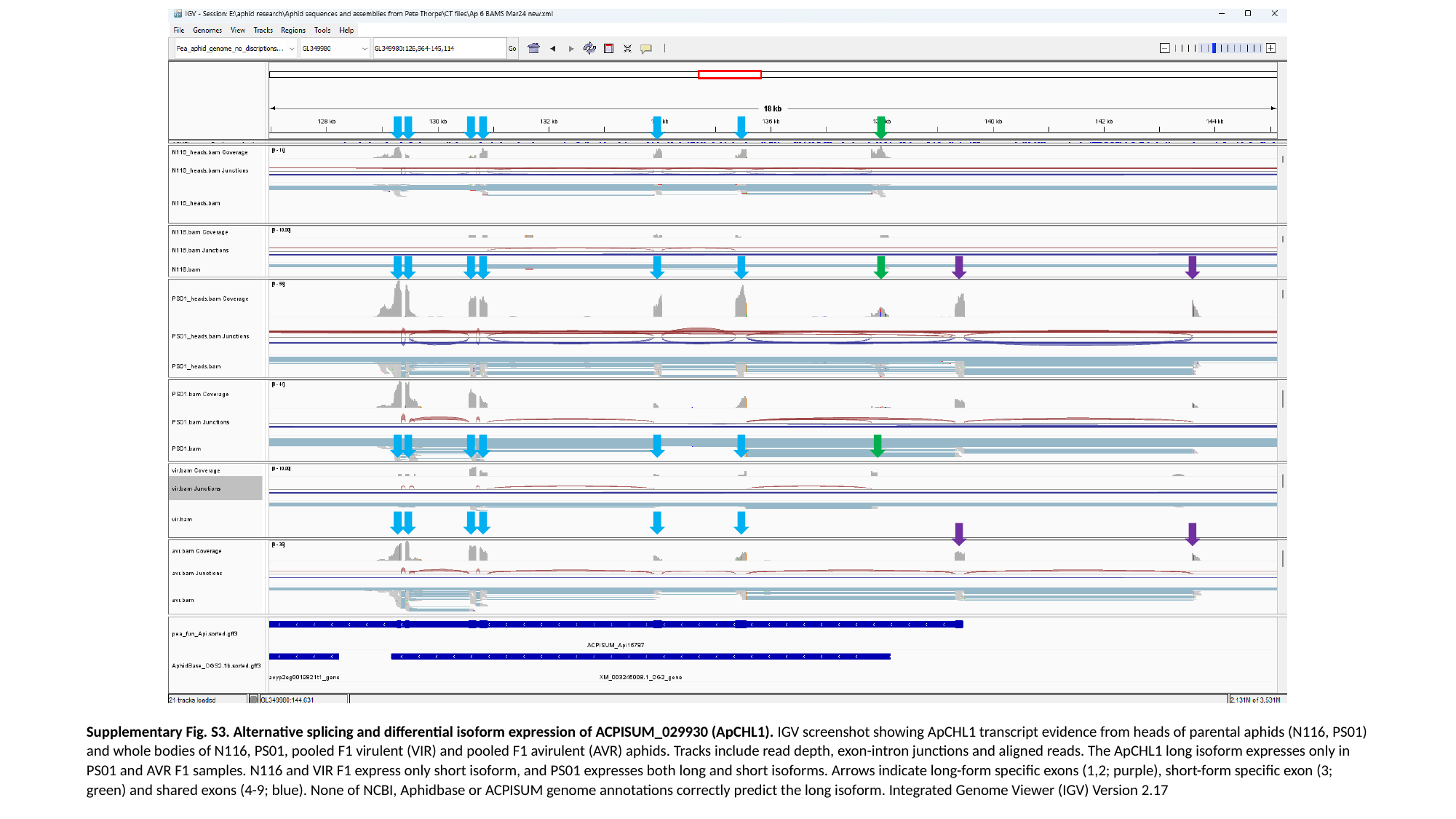

Supplementary Fig. S3. Alternative splicing and differential isoform expression of ACPISUM_029930 (ApCHL1). IGV screenshot showing ApCHL1 transcript evidence from heads of parental aphids (N116, PS01) and whole bodies of N116, PS01, pooled F1 virulent (VIR) and pooled F1 avirulent (AVR) aphids. Tracks include read depth, exon-intron junctions and aligned reads. The ApCHL1 long isoform expresses only in PS01 and AVR F1 samples. N116 and VIR F1 express only short isoform, and PS01 expresses both long and short isoforms. Arrows indicate long-form specific exons (1,2; purple), short-form specific exon (3; green) and shared exons (4-9; blue). None of NCBI, Aphidbase or ACPISUM genome annotations correctly predict the long isoform. Integrated Genome Viewer (IGV) Version 2.17

### Slide 4
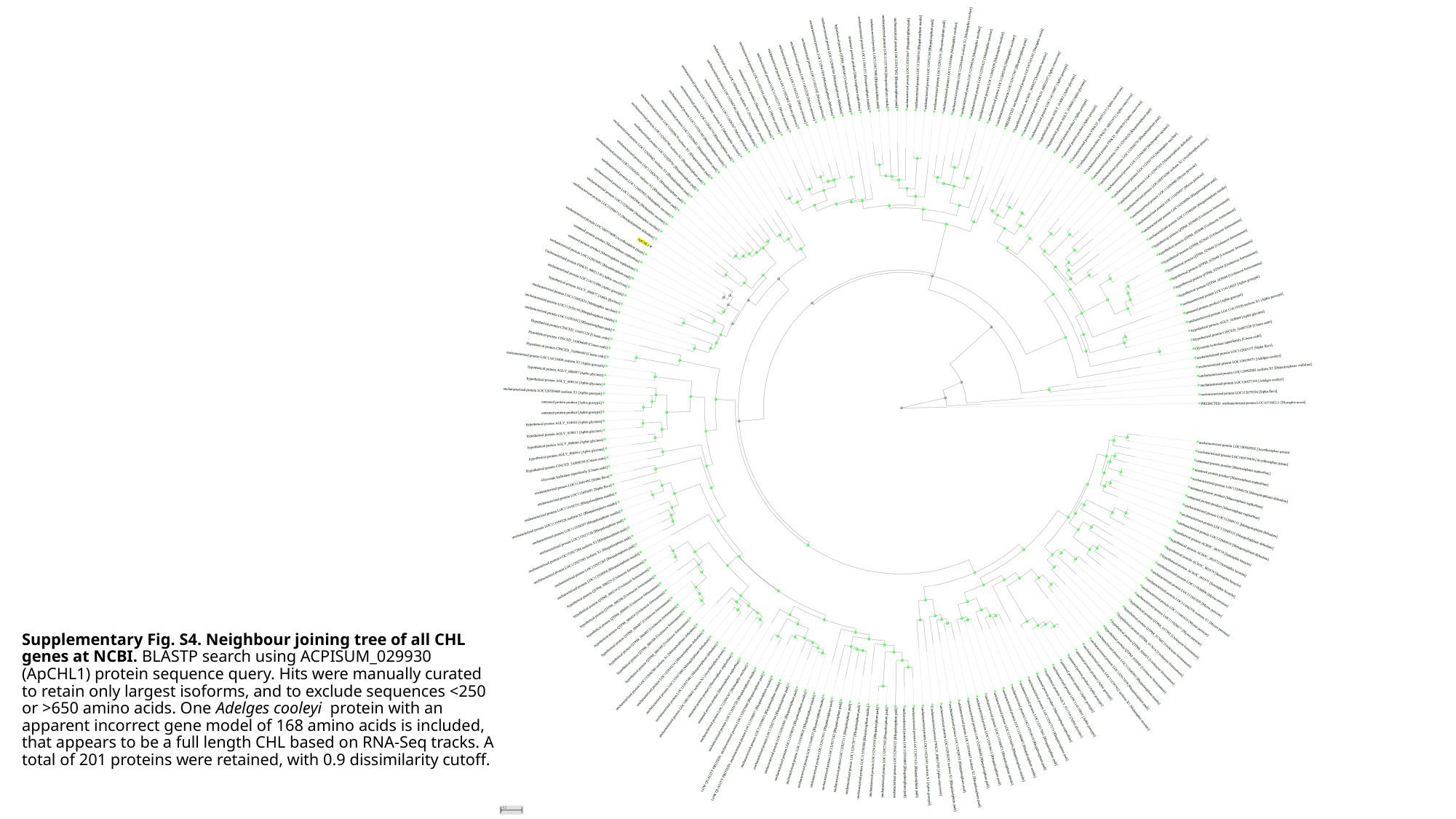

#### Supplementary Fig. S4. Neighbour joining tree of all CHL genes at NCBI. BLASTP search using ACPISUM_029930 (ApCHL1) protein sequence query. Hits were manually curated to retain only largest isoforms, and to exclude sequences <250 or >650 amino acids. One Adelges cooleyi protein with an apparent incorrect gene model of 168 amino acids is included, that appears to be a full length CHL based on RNA-Seq tracks. A total of 201 proteins were retained, with 0.9 dissimilarity cutoff.

### Slide 5
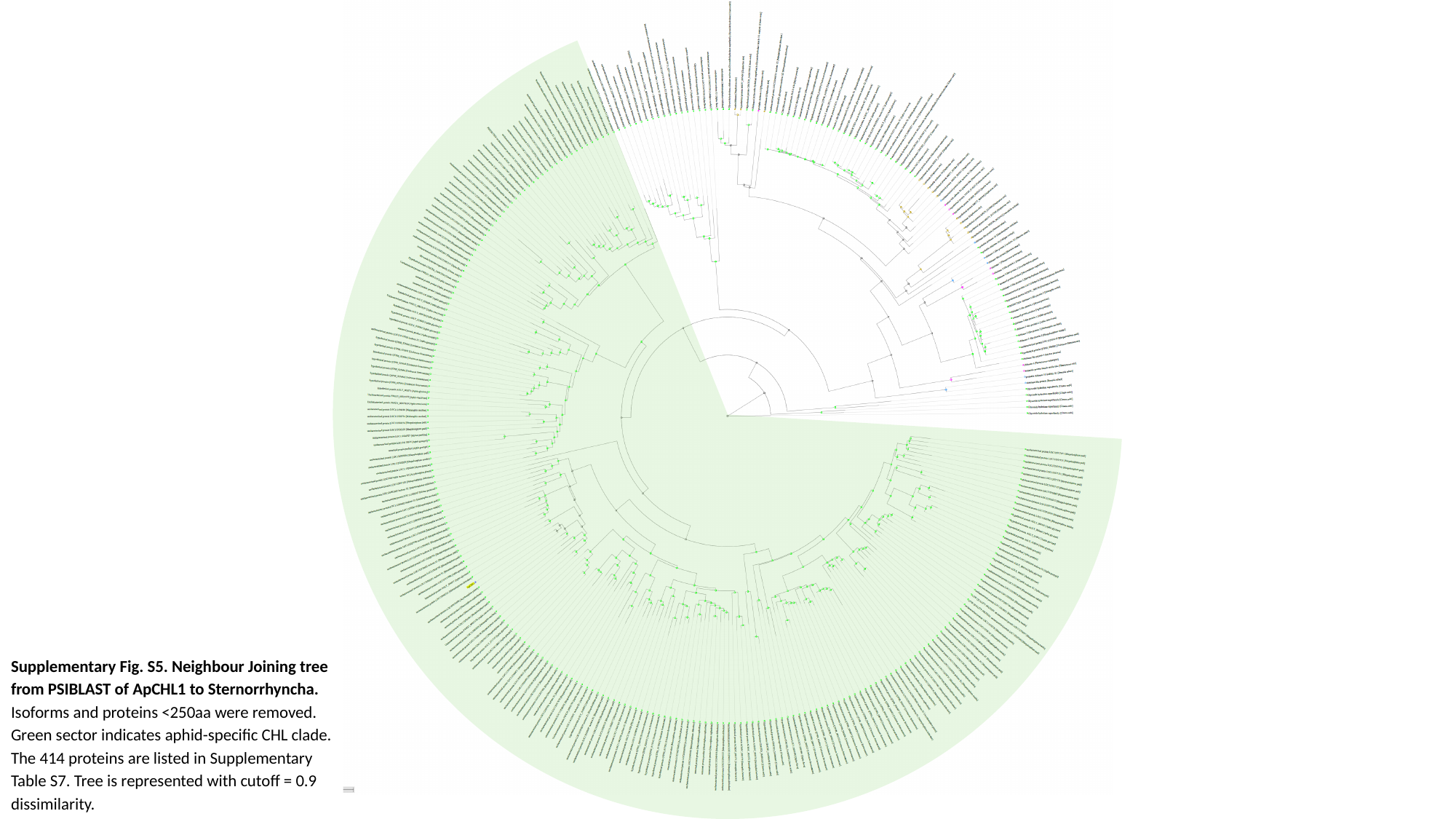

Supplementary Fig. S5. Neighbour Joining tree from PSIBLAST of ApCHL1 to Sternorrhyncha. Isoforms and proteins <250aa were removed. Green sector indicates aphid-specific CHL clade. The 414 proteins are listed in Supplementary Table S7. Tree is represented with cutoff = 0.9 dissimilarity.

### Slide 6
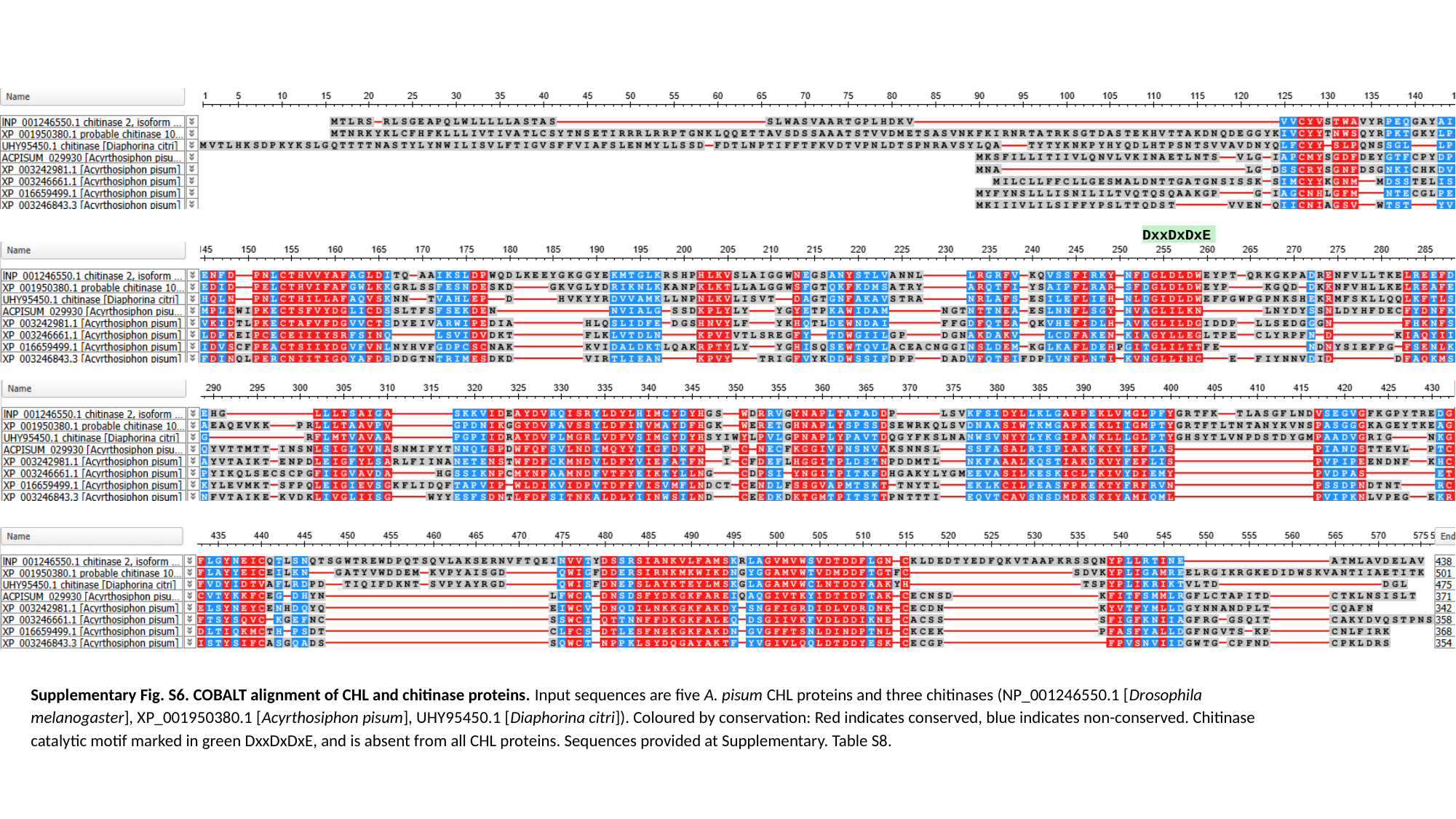

DxxDxDxE
Supplementary Fig. S6. COBALT alignment of CHL and chitinase proteins. Input sequences are five A. pisum CHL proteins and three chitinases (NP_001246550.1 [Drosophila melanogaster], XP_001950380.1 [Acyrthosiphon pisum], UHY95450.1 [Diaphorina citri]). Coloured by conservation: Red indicates conserved, blue indicates non-conserved. Chitinase catalytic motif marked in green DxxDxDxE, and is absent from all CHL proteins. Sequences provided at Supplementary. Table S8.

### Slide 7
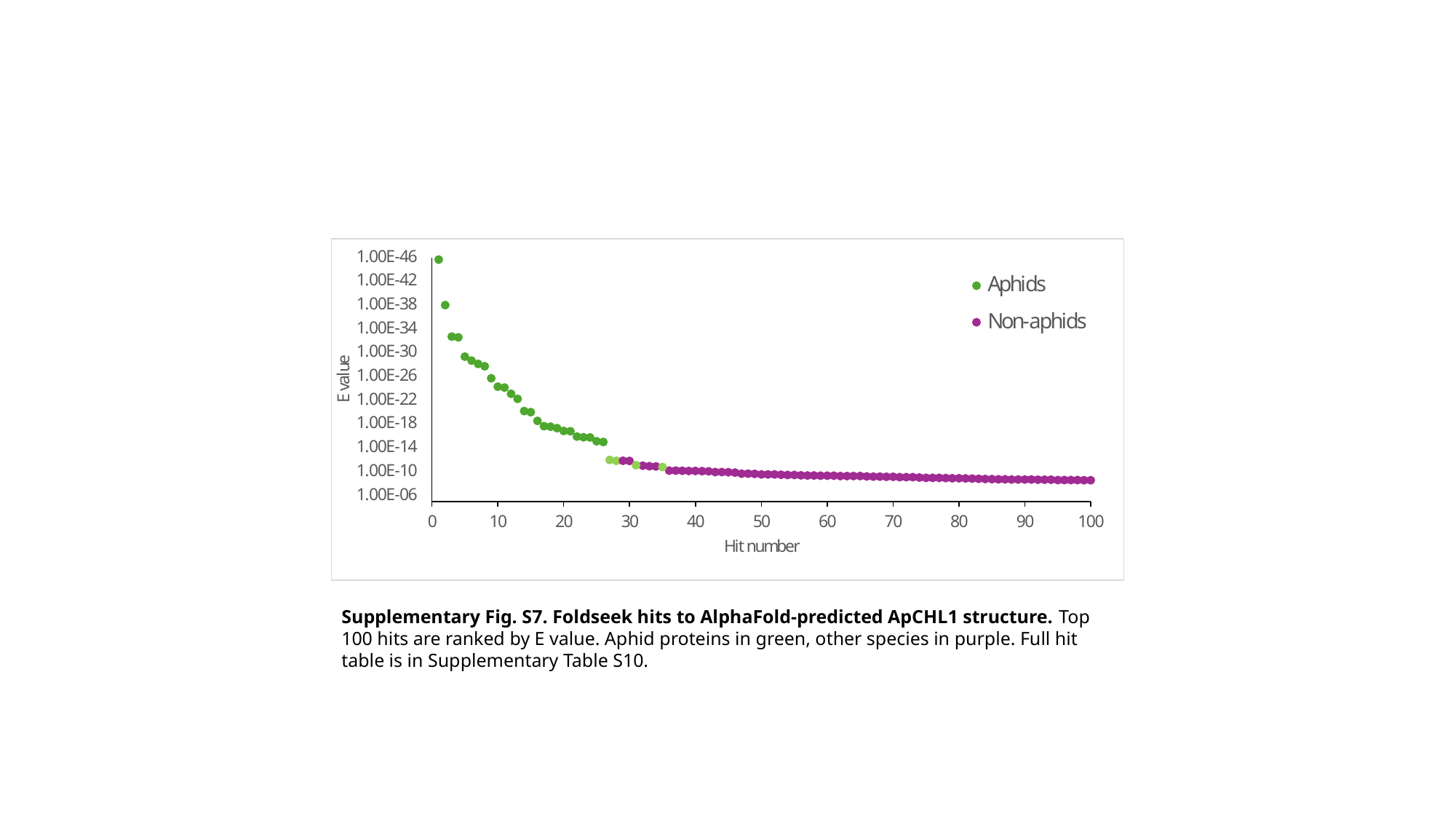

Supplementary Fig. S7. Foldseek hits to AlphaFold-predicted ApCHL1 structure. Top 100 hits are ranked by E value. Aphid proteins in green, other species in purple. Full hit table is in Supplementary Table S10.

### Slide 8
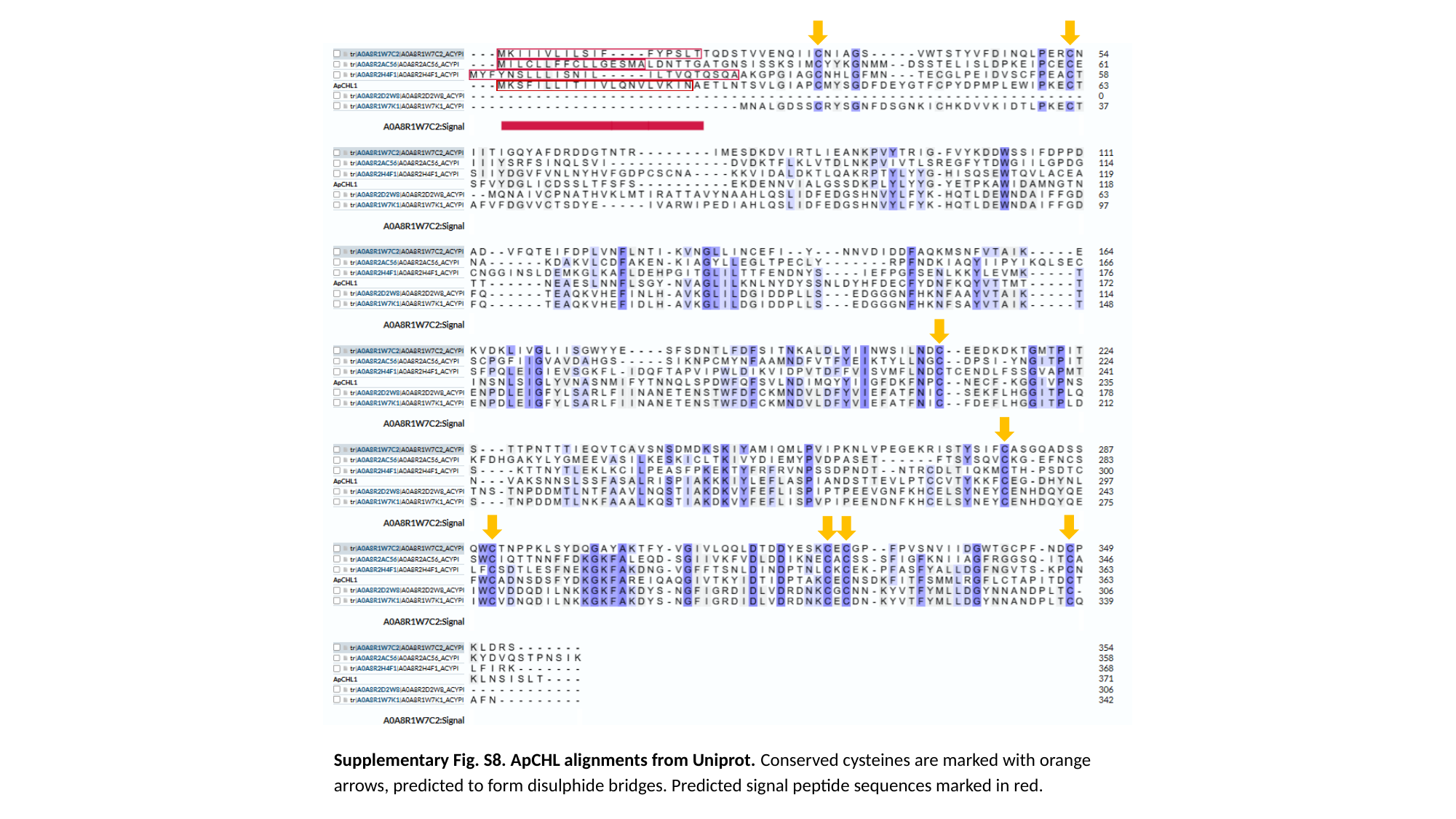

Supplementary Fig. S8. ApCHL alignments from Uniprot. Conserved cysteines are marked with orange arrows, predicted to form disulphide bridges. Predicted signal peptide sequences marked in red.
